# Safb1 regulates cell fate determination in adult neural stem cells by enhancing Drosha cleavage of NFIB mRNA

**DOI:** 10.1101/2021.10.22.465432

**Authors:** Niklas Iffländer, Chiara Rolando, Elli-Anna Balta, Pascal Forcella, Tanzila Mukhtar, Thomas Bock, Verdon Taylor

## Abstract

During brain homeostasis, stem cell fate determination is crucial to guarantee function, adaptation and regeneration while preventing neurodegeneration and cognitive impairment. How neural stem cells (NSCs) are instructed to generate neurons or glia is not well understood. Here we addressed how fate is resolved in multipotent adult hippocampal NSCs, and identify Scaffold Attachment Factor B1 (Safb1) as a determinant of neuron production by blocking glial commitment. Safb1 is sufficient to block oligodendrocytic differentiation of NSCs by preventing expression of the transcription factor NFIB at the post-transcriptional level. Detailed interrogation of the Drosha interactome and functional validation revealed that Safb1 enhances NFIB mRNA cleavage in a Drosha-dependent fashion. Thus, our study provides a cellular mechanism for selective NSC fate regulation by post-transcriptional destabilization of mRNAs. Given the importance of NSC maintenance and fate determination in the adult brain, our findings have major implications for cell-specific gene expression, brain disease and aging.

## Introduction

NSCs are multipotent stem cells and their neuronal and glial fate determination are precisely controlled. NSCs generate new neurons during brain development before switching to generate glia and the subsequent elimination in most regions of the postnatal brain. While neurogenesis is prominent at embryonic stages, in the adult vertebrate brain the generation of new neurons is predominantly restricted to two niches that maintain NSCs, the ventricular subventricular zone (V-SVZ) of the lateral ventricles and the subgranular zone of the hippocampal dentate gyrus (DG) (Goncalves et al., 2016; Obernier and Alvarez-Buylla, 2019). Whereas V-SVZ NSCs generate multiple neuron types, astrocytes and oligodendrocytes, DG NSCs predominantly generate glutamatergic granule neurons and to a less extent astrocytes but not oligodendrocytes (Bonaguidi et al., 2011; Bonzano et al., 2018; Pilz et al., 2018; Seri et al., 2004). Adult hippocampal neurogenesis plays a central role in memory formation, plasticity and learning in rodents, but also in other species including humans (Berg et al., 2019; Boldrini et al., 2018; Eriksson et al., 1998; Gage, 2019; Moreno-Jimenez et al., 2019; Spalding et al., 2013). Although still controversially discussed, hippocampal neurogenesis persists into the late decades of life in humans, and changes in neuron production might be linked to neurodegenerative diseases, including Alzheimer’s Disease (Beckervordersandforth and Rolando, 2019; Boldrini *et al*., 2018; Kempermann et al., 2018; Moreno-Jimenez *et al*., 2019; Sorrells et al., 2018; Tobin et al., 2019). In rodents, adult NSC regulation is fairly well understood at the molecular and physiological levels, however, the influence and importance of post-transcriptional regulation in adult neurogenesis is starting to become clearer (Baser et al., 2019; Pilaz and Silver, 2015).

The ribonuclease Drosha is a core component of the microRNA pathway and forms a trimeric complex with two DGCR8 proteins to generate the microRNA Microprocessor (Han et al., 2004; Nguyen et al., 2015). The Microprocessor processes pri-microRNA stem loop hairpins (HPs) to generate pre-microRNAs in a specific pattern (Kim et al., 2017). Pre-microRNAs are further processed into mature microRNAs by Dicer before being directed to target mRNAs by the RISC complex. However, Drosha is involved in different, non-canonical, post-transcriptional gene regulation mechanisms beyond its primary function in microRNA biogenesis (Chong et al., 2010; Lee and Shin, 2018; Rolando and Taylor, 2017). Although microRNAs are important for terminal neuronal differentiation (Yoo et al., 2011), Drosha maintains the embryonic and hippocampal NSC pool independent of its microRNA biogenesis activity (Knuckles et al., 2012; Rolando et al., 2016). The expression of the proneural gene Neurogenin2 (Ngn2) is tightly controlled by Drosha, which cleaves evolutionary conserved HPs in Ngn2 mRNA transcripts (Knuckles *et al*., 2012). To date, multiple Drosha targeted mRNAs have been identified in NSCs in addition to that of Ngn2, including those of NeuroD1, NeuroD6 and Nuclear Factor I/B (NFIB) (Knuckles *et al*., 2012; Rolando *et al*., 2016). The transcription factor NFIB is required and sufficient to promote glial-fate specification in neural progenitors (Deneen et al., 2006). In the adult hippocampus, Drosha represses NFIB mRNA and prevents DG NSCs acquiring an oligodendrocytic fate (Rolando *et al*., 2016). NFIB mRNA contains two Drosha interacting HPs, located in the 5’ untranslated region (UTR) and 3’ UTR. Despite that both HPs being bound by Drosha, only the NFIB 3’ UTR HP is cleaved (Rolando *et al*., 2016). How the cleavage specificity towards these HPs is achieved has still to be investigated.

Drosha is ubiquitously expressed, and its protein-protein interactions are not restricted to DGCR8. Different Drosha-binding partners have been identified underlining the diversity in Drosha functions on RNA including splicing and transcriptional regulation (Rouillard et al., 2016; Spadotto et al., 2020). However, the composition of the Drosha complexes that regulate NSC maintenance and differentiation in the adult brain are unknown. Thus, it is unclear how Drosha-mediated cleavage of mRNAs is regulated to control NSC pool maintenance and cell fate in what is potentially a cell-type specific manner. We hypothesized that Drosha activity in controlling cell fate by direct mRNA degradation is regulated by associated proteins. Using proteomic analyses, we identified Drosha interacting proteins in DG hippocampal NSCs. We found that the RNA binding protein Safb1 regulates Drosha’s activity in the destabilization of NFIB mRNA. Safb1 is implicated in multiple cellular processes, including cellular stress, DNA damage response, cell growth and apoptosis and has been linked to microRNA biogenesis (Altmeyer et al., 2013; Townson et al., 2000; Treiber et al., 2017). Although Safb1 is expressed in many tissues, its expression is particularly high in the brain (Rivers et al., 2015; Townson et al., 2003). Here we demonstrate that Safb1 levels are high in DG NSCs blocking glial differentiation through regulating Drosha cleavage of the NFIB mRNA, and are lower in V-SVZ NSCs thus allowing their differentiation to an oligodendrocytic fate.

## Results

### Identification of Drosha-binding partners in DG NSCs

In order to examine the Drosha DG NSC interactome and identify proteins that may regulate canonical and non-canonical activities of Drosha in controlling NSC fate, we performed Drosha co-immunoprecipitation (co-IP) followed by tandem mass spectrometry (MS^2^) from adult mouse DG NSCs (**Figure 1A** and **Figure 1-figure supplement 1A,B**). 165 proteins co-precipitated with Drosha (p < 0.05, log_2_ fold change ≥ 3, FDR ≤ 0.001, peptide count ≥ 4; **Figure 1B** and **Table S1**), the majority of which are RBPs (138; 84%) (Gerstberger et al., 2014; Huang et al., 2018). We compared our DG NSC Drosha interactome dataset with a previous Drosha IP performed from human embryonic kidney cells (HEK293T) and found an overlap of 24 proteins (Macias et al., 2015). Comparison of the interactome with the 20 protein components of the large Drosha complex described previously (CORUM protein complexes dataset) revealed an overlap of 11 proteins (**Figure 1B,C**) (Rouillard *et al*., 2016). Therefore, approximately 50% of the canonical large Drosha complex proteins identified in other cellular systems were identified as Drosha partner proteins in the DG NSC interactome (**Figure 1B,C**). However, we identified over 100 novel Drosha partners. As a proof of concept, the canonical Drosha interacting protein and partner in the microRNA Microprocessor complex DGCR8 was highly enriched in the DG NSC Drosha interactome (**Figure 1B**).

**Figure 1:**
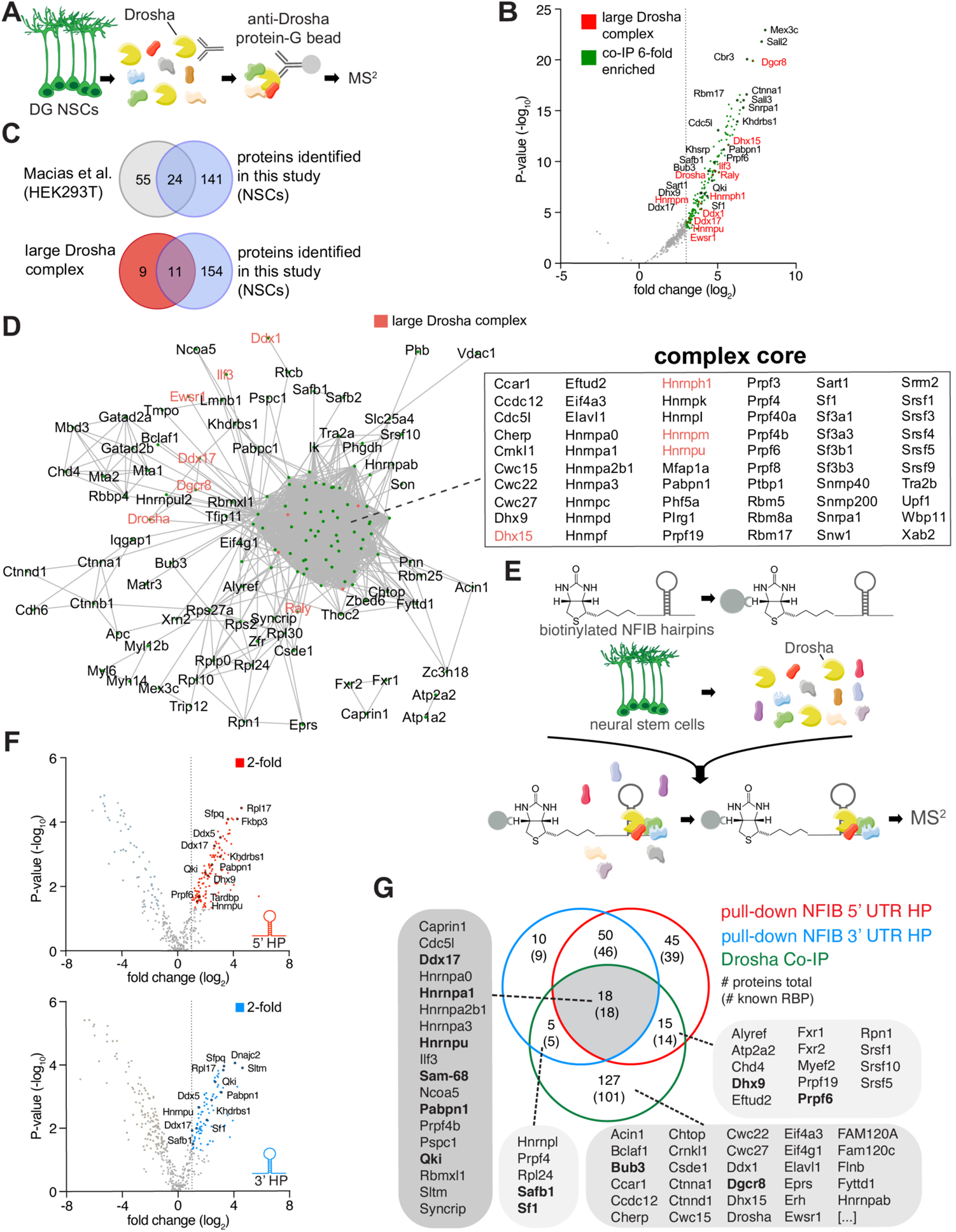
Identification of Drosha and NFIB mRNA with RNA binding proteins in NSCs. **A.** Scheme of the endogenous Drosha co-immunoprecipitation procedure from DG NSCs followed by MS^2^ analysis. **B.** Volcano plot of MS^2^ quantified proteins displaying as fold change log_2_ of Drosha co-precipitated proteins over control (x-axis) and P-value −log_10_ (y-axis). Significantly enriched Drosha associated proteins (green dots) were determined as having P-value < 0.05, log_2_ fold change ≥ 3 and false discovery rate (FDR) *≤* 0.001. Eleven proteins of the known “large Drosha complex” (red) were also enriched in the coprecipitation with Drosha from DG NSCs. Selected novel proteins are also shown (black). For full list of MS^2^ quantified proteins see Table S1. **C.** Venn-diagrams of pairwise comparisons of Drosha interacting proteins identified in DG NSCs in this study and Drosha interacting proteins in HEK293T cells, and the CORUM large Drosha complex (Macias *et al*., 2015; Rouillard *et al*., 2016). **D.** STRING network analysis of Drosha interacting proteins in DG NSCs. Nodes are indicated as green dots, edges correspond to known interactions substantiated by experimental data. Only nodes with one or more edges are displayed, protein isoforms are shown collectively. The proteins in the densely-packed, core complex are listed alphabetically. **E.** Scheme of the NFIB hairpin pull-down assay using biotinylated RNA probes as bait to capture binding proteins followed by precipitation with streptavidin-coupled beads and MS^2^ analysis. **F.** Volcano plots of MS^2^ quantified NFIB 5’ UTR HP (top - red) and NFIB 3’ UTR HP (bottom - blue) interacting proteins displayed as fold change log_2_ (x-axis) versus P-value −log_10_ (y-axis). Significant enriched proteins (red and blue dots) were defined as having P-value < 0.05, log_2_ fold change ≥ 1 and FDR *≤* 0.1. Selected novel proteins are also shown (black). For full list of interacting proteins see Table S1. **G.** Venn-diagram of comparisons between Drosha interacting proteins in DG NSCs (green circle), NFIB 3’ UTR HP binding proteins (blue circle) and NFIB 5’ UTR HP binding proteins (red circle). The numbers indicate the total number of proteins, the numbers in brackets indicate how many of the proteins are known RBPs. Proteins selected for the functional assay are highlighted in the lists in bold type.

**Figure 1-figure supplement 1:**
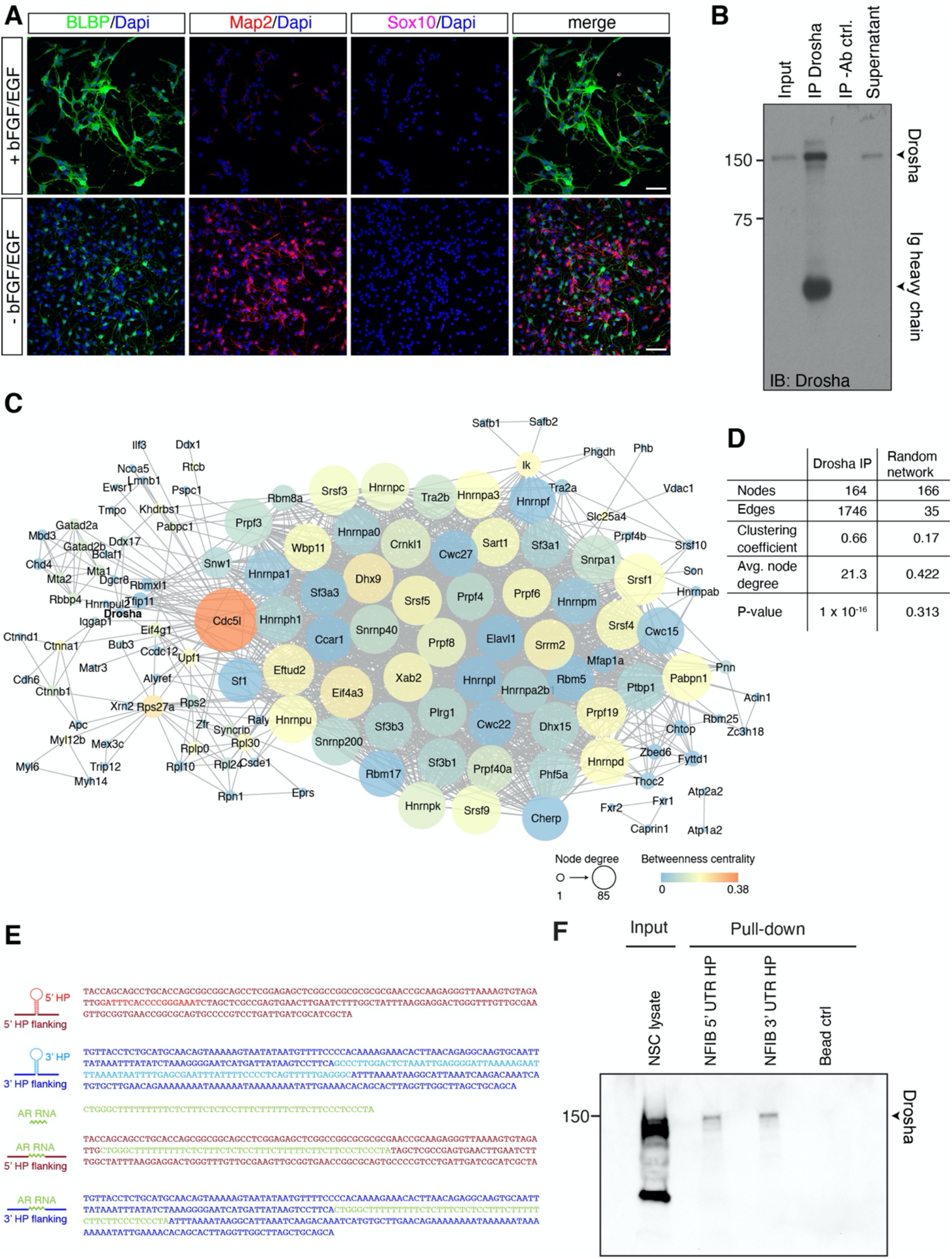
DG NSCs contain large Drosha complexes. **A.** Characterization of DG NSCs under expansion (+bFGF/EGF) and differentiation (-bFGF/EGF) culture conditions. Immunohistochemistry for the progenitor marker BLBP, neuronal protein Map2 and oligodendrocyte protein Sox10. Scale bar 50 µm. **B.** Immunoblot (IB) validation of Drosha immunoprecipitation (IP). IP with anti-Drosha antibodies shows precipitation of Drosha. Negative control precipitation without antibody (-Ab). Input is 2.5% of total lysate used in the IP. Drosha protein and immunoglobulin (Ig) heavy chain from the precipitating antibody are indicated. **C.** STRING network analysis of the Drosha interacting proteins identified by MS^2^ (related to Figure 1D). Node size corresponds to node degree, node color corresponds to betweenness centrality, edges exclusively correspond to known interactions based on experimental data and databases. Only nodes with one or more edges are displayed, protein isoforms were analyzed collectively. **D.** Common network parameters for Drosha IP network compared with a random network of similar node size. (Related to Figure 1D). **E.** Base sequence of the NFIB 5’ UTR HP and 3’ UTR HP RNA probes, the binding sequence for HuR from the androgen receptor mRNA (AR), and AR RNA-NFIB UTR hybrid probes used in the pull-down experiments. The colors correspond to the relative RNA domains shown in the secondary structure schemes. **F.** Immunoblot (IB) validation of Drosha precipitation with NFIB RNA 3’ UTR HP and NFIB 5’ UTR HP pull-down probes. Negative control precipitation is bead-only control (Bead ctrl).

**Figure 1-figure supplement 2:**
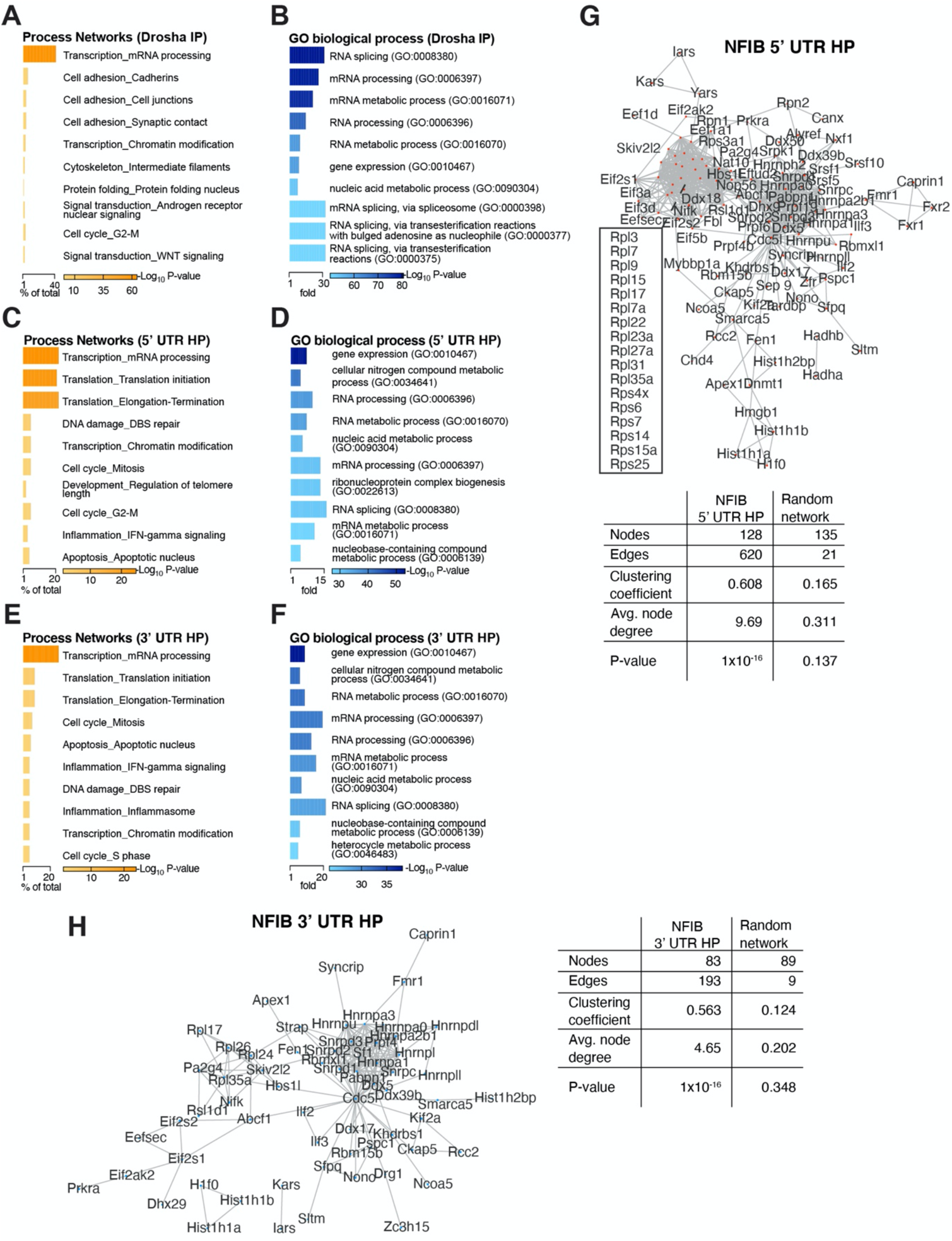
Process analysis of Drosha and NFIB mRNA interacting proteins. **A.** MetaCore enrichment analysis of process networks for the identified Drosha-interacting proteins. Bar length corresponds to percentage of protein number out of total identified Drosha interacting proteins for each of the indicated categories. P-values are indicated by color. **B.** GO terms (PANTHER) of biological processes for the identified Drosha interacting proteins. Bar length corresponds to fold change enrichment per category. P-values are indicated by color. **C, D.** MetaCore enrichment analysis of Process Networks and GO terms (PANTHER) of biological processes for the NFIB 5’ UTR HP interacting proteins. Bar length corresponds to the percentage of protein out of total identified NFIB 5’ UTR HP interacting proteins per category (Process Networks) or to fold change per category (GO biological processes). P-values are indicated by color. **E, F.** MetaCore enrichment analysis of Process Networks and GO terms (PANTHER) of biological processes for the NFIB 3’ UTR HP interacting proteins. Bar length corresponds to the percentage of proteins out of total identified NFIB 3’ UTR HP interacting proteins per category (Process Networks) or to fold change per category (GO biological processes). P-values are indicated by color. **G, H.** STRING network analysis of NFIB 5’ UTR HP and 3’ UTR HP interacting proteins. Nodes are indicated by colored dots, edges exclusively correspond to known interactions from experimental data and databases. Only nodes with one or more edges are displayed, protein isoforms are shown collectively. Tables: Common network parameters for pull-down networks compared with random networks of similar node size for the NFIB 5’ UTR HP (left) and NFIB 3’ UTR HP (right) probes.

**Figure 1-figure supplement 3:**
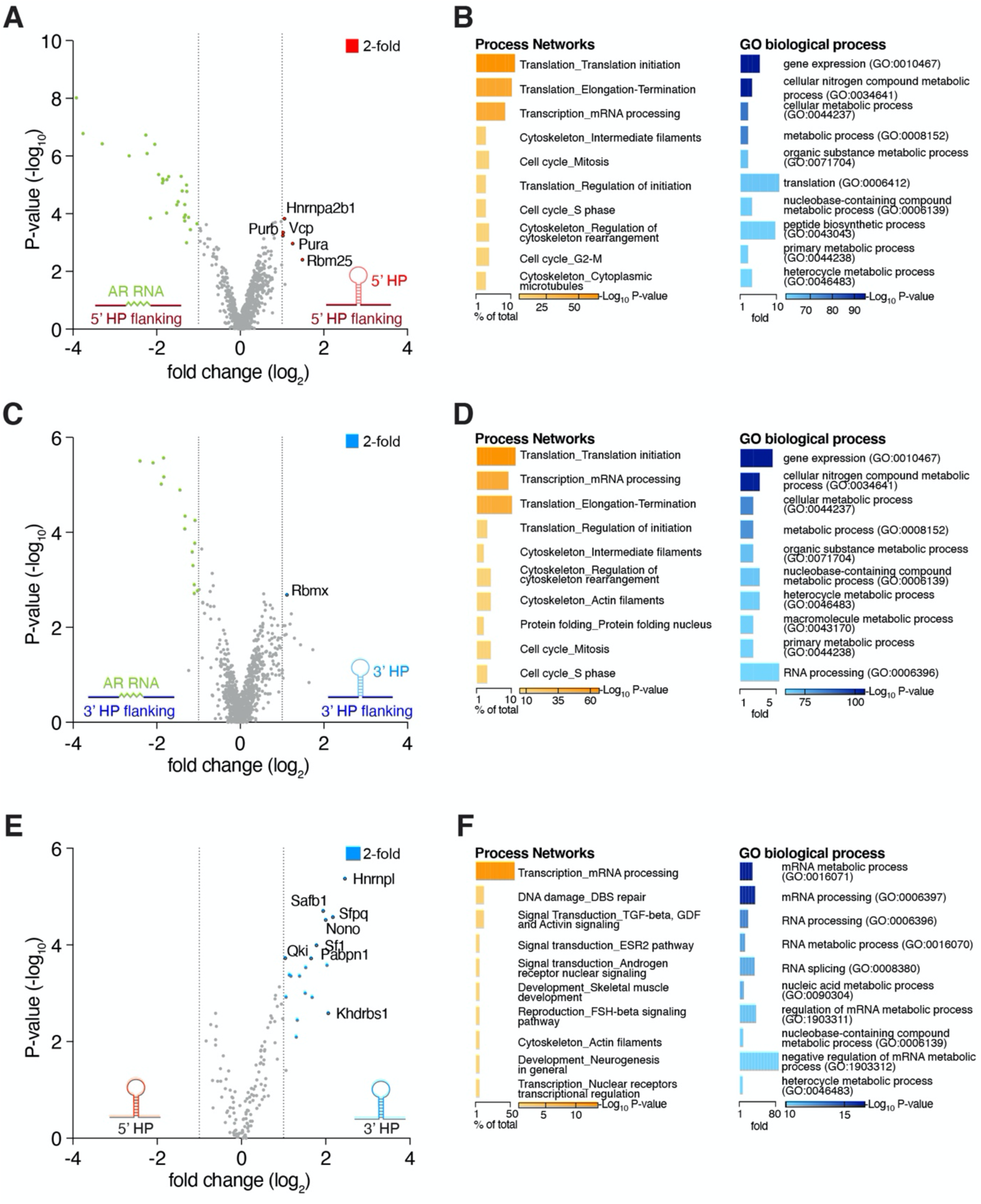
NFIB mRNA interactome analysis uncovers regional binding preferences. **A.** Proteomic analysis of interactors with the NFIB 5’ UTR HP (including the HP flanking sequences). Volcano plot of MS^2^ quantified proteins displaying log_2_ fold change (x-axis) versus −log_10_ P-value (y-axis). **B.** NFIB 5’ UTR HP flanking region protein association analysis. The dataset of flanking proteins includes proteins not enriched for 5’ UTR HP or control in the MS^2^ analysis. Non-specific bead binding proteins were subtracted from the analysis. MetaCore enrichment analysis of process networks and GO terms (PANTHER) of biological processes for identified NFIB 5’ UTR HP flanking binding proteins. **C.** Proteomics analysis of interactors with the NFIB 3’ UTR HP (including the HP flanking sequences). Volcano plot of MS^2^ quantified proteins displaying log_2_ fold change (x-axis) versus −log_10_ P-value (y-axis). **D.** NFIB 3’ UTR HP flanking region protein association analysis. The dataset of flanking proteins includes proteins not enriched for 3’ UTR HP or control in MS^2^ analysis. Non-specific bead binding proteins were subtracted from the analysis. MetaCore enrichment analysis of process networks and GO terms (PANTHER) of biological processes for identified NFIB 3’ UTR HP flanking binding proteins. **E.** Proteomics analysis of NFIB 3’ UTR HP versus NFIB 5’ UTR HP interactors. Volcano plot of MS^2^ quantified proteins displaying log_2_ fold change (x-axis) versus −log_10_ P-value (y-axis). **F.** MetaCore enrichment analysis of process networks and GO terms (PANTHER) of biological processes for specific NFIB 3’ UTR HP flanking proteins. Volcano plots: Significant enrichment for highlighted proteins (colored dots) was achieved by P-value < 0.05, log_2_ fold change ≥ 1 and FDR *≤* 0.1. A subset of proteins is displayed. Bar plots: Bar length corresponds to percentage of protein number out of total identified NFIB UTR HP interacting proteins per category or to fold change per category, respectively. P-values indicated by color. STRING functional protein association network analysis within the DG NSC Drosha interactome (considering only experimentally determined data and curated databases) (Szklarczyk et al., 2019) revealed one major complex indicating the close connectivity between interactors (Figure 1D and Figure 1-figure supplement 1C,D). Strikingly, many of the known Drosha interactors, including Drosha itself, were not positioned in the core of the complex indicating additional mediators between Drosha and distant co-interactors. In summary, we identified novel Drosha binding partners in adult DG NSCs, many of which are involved in transcriptional regulation and RNA biogenesis.

Process network analysis (MetaCore) indicated that 56 of the 165 Drosha interacting proteins (34%, p = 10^−61^) are involved in transcriptional regulation and mRNA processing (**Figure 1-figure supplement 2A**). Conversely, only 2 - 7% (3 - 11) of the Drosha interacting proteins have been linked to other process networks and these include translation, cell cycle, cell adhesion and the cytoskeleton (p = 10^−1^ − 10^−3^) (**Figure 1-figure supplement 2A**). In order to gain insights into the biological functions of the Drosha interactors, we performed Gene Ontology (GO) enrichment analysis of biological processes. The top GO term of the Drosha interactors in NSCs was RNA splicing (GO:0008380) with >30-fold enrichment, followed by RNA processing (**Figure 1-figure supplement 2B**). As Drosha is primarily linked to RNA regulation, translation and microRNAs, these findings support the specificity of the endogenous Drosha pull-down assay.

**Figure 2:**
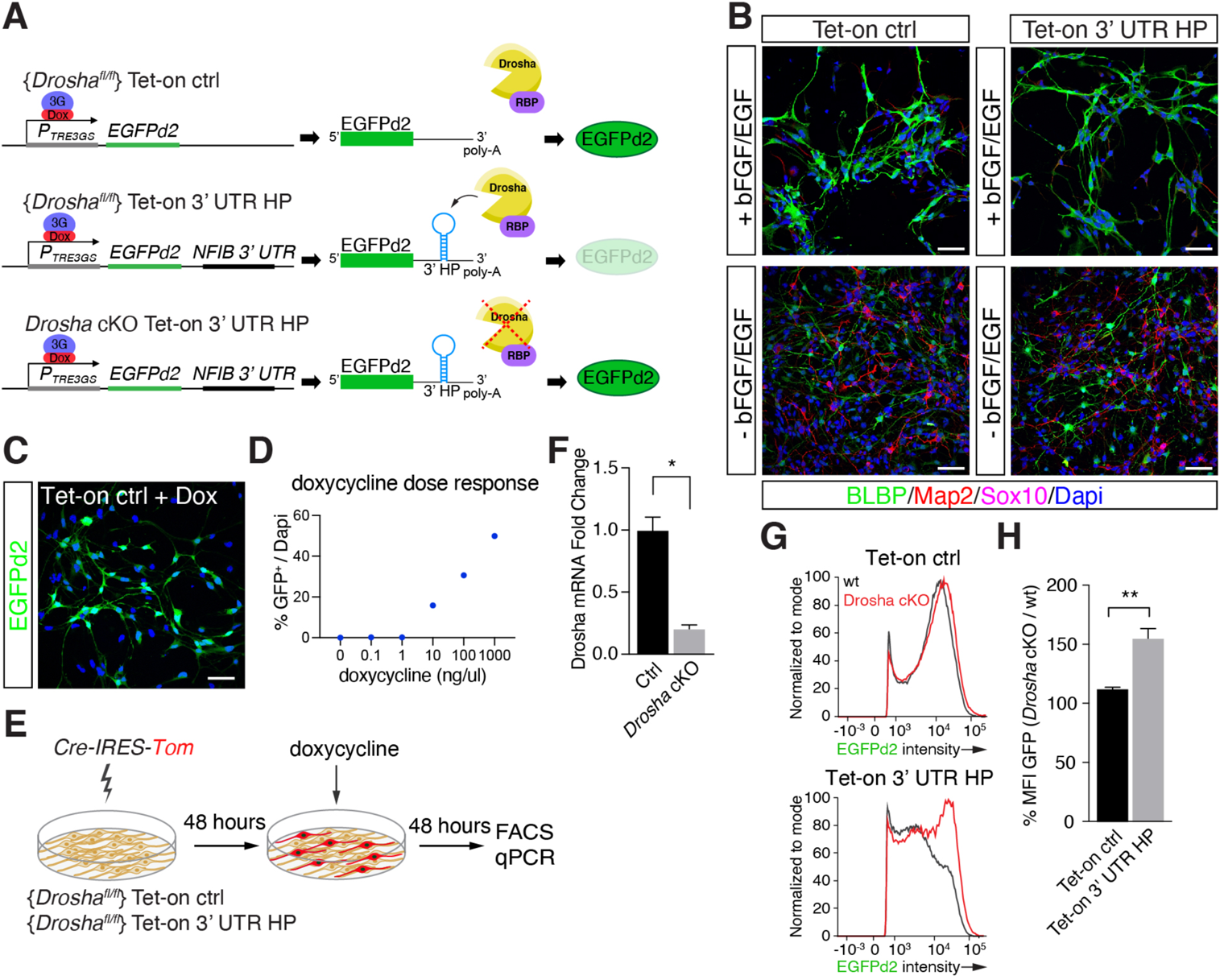
Conditional DG NSCs report Drosha processing of NFIB constructs. **A.** Scheme of the experimental paradigm using the Tet-on reporter lines to examine the effects of the NFIB 3’ UTR HP (composed of NFIB 3’ UTR HP inserted into the untranslated region downstream of destabilized EGFP (EGFPd2)) on expression. Stable floxed *Drosha* (*Drosha^fl/fl^*) DG NSCs lines carrying doxycycline inducible Tet-on ctrl, or Tet-on 3’ UTR HP constructs were generated and selected. Drosha/RBP complexes regulate stability of the reporter RNA and EGFPd2 expression levels. Deletion of Drosha stabilizes the Tet-on 3’ UTR HP construct mRNA and EGFPd2 expression. **B.** Characterization of the Tet-on ctrl and NFIB 3’ UTR HP (Tet-on 3’ UTR) expressing DG NSCs under expansion (+bFGF/EGF) and differentiation (-bFGF/EGF) culture conditions. Immunohistochemistry for the progenitor marker BLBP, neuronal protein Map2 and oligodendrocyte protein Sox10. Scale bars 50 µm. **C.** Expression of EGFPd2 by doxycycline-induced (48 hours) Tet-on ctrl DG NSC line (Tet-on ctrl). Scale bar 50 µm. **D.** Doxycycline dose response curve of Tet-on ctrl DG NSC line. Quantification of EGFPd2^+^ (GFP^+^) cells over total cells (DAPI). **E.** Experimental paradigm for *Drosha* conditional deletion (*Drosha* cKO) experiments from Tet-on ctrl and Tet-on 3’ UTR HP DG NSCs followed by quantitative FACS analysis for EGFPd2 expression and RT-qPCR. **F.** RT-qPCR quantification of Drosha mRNA levels before (Ctrl) and after *Drosha* cKO from Tet-on 3’ UTR HP DG NSCs. n = 4, two-tailed Mann-Whitney test: *p<0.05. Error bars SEM. **G.** FACS analysis of EGFPd2 fluorescence by Tet-on ctrl and Tet-on 3’ UTR HP DG NSCs after doxycycline induction (48 hours). EGFPd2 intensity (x-axis) versus cell number normalized to mode (y-axis) of Tet-on ctrl and Tet-on 3’ UTR HP DG NSC lines before *Drosha* cKO (wt: black line) and after *Drosha* cKO (red line). Note the recovery of high EGFPd2 expressing cells (intensity >10^4^) in the *Drosha* cKO Tet-on 3’ UTR HP DG NSC (red line) compared to the same cells before Drosha deletion (wt: black line). **H.** Quantification of median fluorescence intensity of EGFPd2 (GFP) from *Drosha* cKO over wt in Tet-on ctrl and Tet-on 3’ UTR HP lines. n = 5, two-tailed Mann-Whitney test: **p<0.01. Error bars SEM.

### Identification of interactors with NFIB 5’ UTR and 3’ UTR HP RNA sequences

With the compendium of Drosha associated proteins in hand, we asked how these may influence Drosha activity to enable non-canonical destabilization of specific mRNAs and influence NSC fate. NFIB is required and sufficient to induce glial-fate specification of NSCs, and necessary for hippocampal development and myelination. NFIB expression is repressed in DG NSCs by Drosha-mediated post-transcriptional cleavage of its mRNA (Deneen *et al*., 2006; Rolando *et al*., 2016). The 5’ UTR and 3’ UTR of the NFIB mRNA contain evolutionary conserved HPs that are bound by Drosha (Rolando *et al*., 2016). To determine the Drosha complexes that potentially control NFIB mRNA stability, we performed RNA pull-down experiments using the critical NFIB mRNA regulatory regions as bait (Rolando *et al*., 2016).

We biotinylated the transcripts of the NFIB HPs including flanking sequences and pulled-down proteins from DG NSC lysates, analyzing the associated protein complexes by MS^2^ (**Figure 1E** and **Figure 1-figure supplement 1E**). As a proof of concept, we confirmed that Drosha was precipitated with both the NFIB 5’ UTR and 3’ UTR HPs (**Figure 1-figure supplement 1F**). We identified 128 and 83 proteins that bind the NFIB 5’ UTR and 3’ UTR HPs, respectively (p < 0.05, log_2_ fold change ≥ 1, FDR *≤* 0.1, peptide count ≥ 4), and that were significantly enriched compared to pull-downs with the proximal 3’ UTR of the androgen receptor (AR) mRNA that was used as a control bait (**Figure 1F** and **Table S2**).

The majority of the NFIB HP interactors (117 of the 128 5’ UTR HP bound proteins and 78 of the 83 3’ UTR HP bound proteins) are known RBPs (Gerstberger *et al*., 2014; Huang *et al*., 2018). MetaCore and GO term analysis of process networks revealed that 22% and 27% of the NFIB 5’ UTR HP and 3’ UTR HP binding proteins, respectively, are known to be involved in transcription (**Figure 1-figure supplement 2C-F**). Strikingly, translation initiation (21% of total, P-value 10^−23^) and elongation (22% of total, P-value 10^−21^) were strongly enriched in the 5’ UTR HP bound proteins, but were far less relevant for the 3’ UTR HP interacting proteins (9% of total, P-value 10^−5^ and 10^−4^). Similarly, STRING network analysis of 5’ UTR and 3’ UTR HP bound proteins resulted in comparable findings (**Figure 1-figure supplement 2G,H**). While both interaction datasets include many heterogeneous nuclear ribonucleoproteins (hnRNPs), only the 5’ UTR HP interacting proteins contained an additional complex that comprised many ribosomal proteins (**Figure 1-figure supplement 2G**).

### NFIB interacting RBPs bind the HP flanking regions

The binding of Drosha and its regulatory complexes to cleavage sites in its target mRNAs is not understood (Rolando *et al*., 2016). We assessed whether the proteins that associated with the Drosha target regions on NFIB mRNA bind the HPs directly or the HP flanking sequences in the NFIB 5’ UTR and 3’ UTR HPs. We compared the proteins that bound to the NFIB 5’ UTR HP and 3’ UTR HP probes with those that bound to hybrid RNA controls (AR RNA / NFIB 5’ UTR flanking and AR RNA / NFIB 3’ UTR flanking) where the NFIB HP forming regions had been replaced with the AR control sequence but retained the NFIB 5’ UTR and 3’ UTR HP flanking regions, respectively (**Figure 1-figure supplement 1E** and **Figure 1-figure supplement 3A,C**). Hnrnpa2b1, Purb, Vcp, Pura and Rbm25 were enriched as NFIB 5’ UTR HP interactors whereas Rbmx was the only protein that selectively interacted with the NFIB 3’ UTR HP (**Figure 1-figure supplement 3A,C** and **Table S2**). Thus, the majority of the RBPs in the NFIB 5’ UTR HP and 3’ UTR HP interaction datasets bind to the regions flanking the HPs in the 5’ and 3’ UTRs and not the HPs of the NFIB mRNA. GO biological process analysis revealed that translation (GO:0006412) was 10-fold enriched in the 5’ UTR HP flanking dataset compared to the random control prediction, whereas regulation of gene expression and RNA processing were most enriched in the 3’ UTR HP flanking region binding protein dataset (**Figure 1-figure supplement 3B,D**).

**Figure 3:**
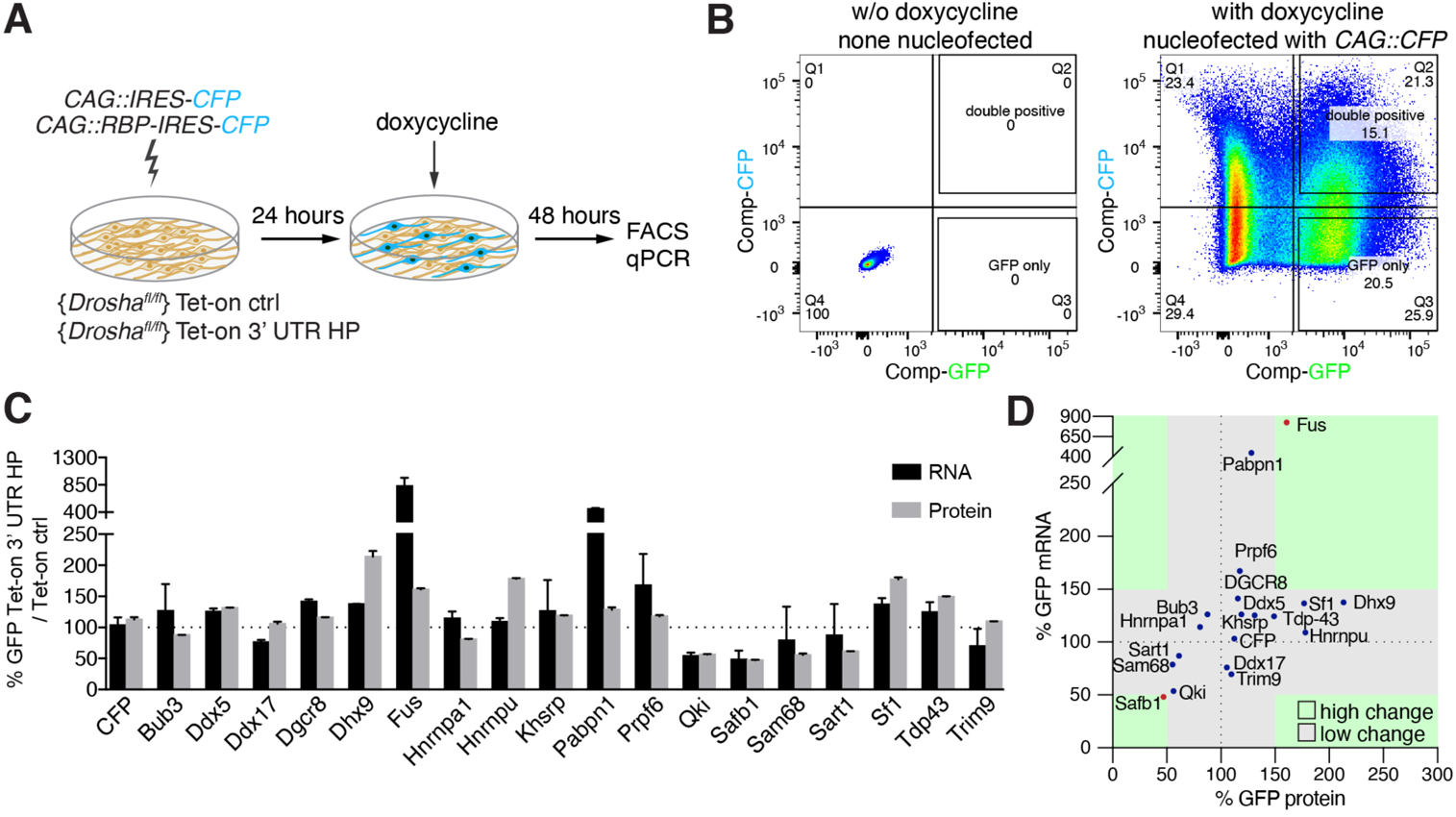
Overexpression of Drosha interactors affects cleavage on NFIB mRNA. **A.** Scheme of the experimental setup to screen the effects of RBPs on the expression of Tet-on ctrl and Tet-on 3’ UTR HP constructs in DG NSCs by FACS and RT-qPCR analysis. RBPs (*CAG::RBP-IRES-CFP*) were expressed in Tet-on ctrl and Tet-on 3’ UTR HP DG NSCs by nucleofection and the levels of EGFPd2 (GFP) protein and mRNA compared to the expression of NSCs expressing a CFP control construct (*CAG::IRES-CFP*) after 48-hours of doxycycline induction. **B.** Representative FACS plot from flow cytometric analyses of Tet-on ctrl DG NSCs with or without nucleofection with the control (*CAG::IRES-CFP*) expression vector, with or without doxycycline induction (48 hours). Untransfected and uninduced Tet-on ctrl DG NSCs do not show EGFPd2 (GFP) or CFP expression compared to doxycycline-induced and *CAG::IRES-CFP* nucleofected Tet-on ctrl DG NSCs. **C.** Analysis of RBP effects on Tet-on 3’ UTR HP construct expression in doxycycline-induced Tet-on 3’ UTR HP DG NSCs. Nucleofected cells were sorted by FACS, gating on the CFP^+^ cells. EGFPd2 (GFP) levels of CFP^+^ cells were quantified by flow cytometry and RNA isolated for RT-qPCR analysis. Quantification of relative EGFPd2 (GFP) mRNA (RT-qPCR) and protein levels (flow cytometry) following overexpression of RPBs by nucleofection in Tet-on 3’ UTR HP DG NSCs compared to control vector (*CAG::IRES-CFP*) transfected cells. To eliminate effects of the RBPs on transcription, RNA stability or translation not linked to the NFIB 3’ UTR HP, the changes in expression were calculated as the differences in EGFPd2 levels in Tet-on 3’ UTR HP DG NSCs and Tet-on ctrl DG NSCs (%GFP Tet-on 3’ UTR HP / Tet-on ctrl). Black dotted line indicates no change. Error bars SEM. **D.** Summary of RBP effects on Tet-on 3’ UTR HP relative to Tet-on ctrl construct expression in DG NSCs. Relative EGFPd2 (GFP) protein levels (fluorescence intensity by FACS, x-axis) and mRNA levels (RT-qPCR, y-axis). Green areas represent changes of +/− >50% of Tet-on 3’ UTR HP relative to Tet-on ctrl construct.

We had found that the 3’ UTR HP of NFIB mRNA is cleaved by Drosha and this contributes to the destabilization of the RNA and blockade of NFIB expression (Rolando *et al*., 2016). Direct comparison of the HP interactors we had previously identified, revealed 18 proteins (p < 0.05, log_2_ fold change ≥ 1, FDR *≤* 0.1, peptide count ≥ 4) that preferentially bound to the NFIB 3’ UTR HP and not to the NFIB 5’ UTR HP, with the highest enrichment for Hnrnpl, Safb1, Sfpq and Nono (**Figure 1-figure supplement 3E**, blue dots and **Table S2**). GO analysis of the NFIB 3’ UTR HP specific interacting proteins showed significant enrichment in negative regulation of mRNA metabolic processes (GO:1903312), consistent with 3’ UTR-mediated mRNA stabilization (**Figure 1-figure supplement 3F**).

We hypothesized that the proteins interacting with Drosha and NFIB HPs could facilitate the Drosha-mediated direct cleavage of target mRNAs. We compared the lists of Drosha, NFIB 3’ and 5’ UTR interacting proteins (**Figure 1G** and **Table S3**) and identified 18 candidates, all of which have previously been reported to be RBPs (Gerstberger *et al*., 2014; Huang *et al*., 2018). As a proof of principle, this subset of 18 RBPs included the Hnrnp family members Hnrnpa1, Hnrnpa2b1 and Hnrnpu, which are known Drosha partners (Macias *et al*., 2015; Rouillard *et al*., 2016). In addition, we identified 5 RBPs that interacted only with Drosha and the 3’ UTR HP region of NFIB and 14 RBPs that bound only to Drosha and the 5’ UTR HP but not the 3’ UTR HP regions of NFIB. The majority of the Drosha-binding partners (127 out of 165 identified proteins in the Drosha IP), including the main partner of Drosha in microRNA biogenesis DGCR8, did not interact with the NFIB 3’ UTR HP or 5’ UTR HP regions indicating that different Drosha complexes are likely involved in different canonical and non-canonical pathways.

### Identification of modulators of non-canonical Drosha activity

To elucidate regulatory functions of the NFIB mRNA-interacting RBPs in Drosha-mediated control of NFIB mRNA stability, we developed a Drosha-cleavage activity reporter system in DG NSCs. We generated stable DG NSC lines carrying a destabilized EGFP (EGFPd2) driven from a doxycycline-regulated expression cassette, and floxed Drosha alleles (*Drosha^fl/fl^*) (Li et al., 1998). One *Drosha^fl/fl^* DG NSC line stably expressed the control EGFPd2 reporter (Tet-on ctrl) (**Figure 2A**), another the same doxycycline-regulated EGFPd2 expression cassette where the NFIB 3’ UTR HP sequence had been inserted downstream of the EGFPd2 coding region (Tet-on 3’ UTR) thereby mimicking the endogenous Drosha-mediated NFIB mRNA processing site (**Figure 2A,B**). Both Tet-on ctrl and Tet-on 3’ UTR HP NSC lines retained stem cell properties including the capacity to generate neurons upon differentiation (**Figure 2B**).

EGFPd2 (referred to hereafter as GFP) expression was induced to submaximal levels by titered administration of doxycycline (**Figure 2C,D** and **3B** (left)). To test whether these inducible DG NSC systems reproduce Drosha-mediated mRNA cleavage of the NFIB mRNA, we analyzed GFP expression following conditional *Drosha* ablation (cKO), induced by transient expression of Cre-recombinase (Cre-IRES-Tom) (**Figure 2A,E,F**) (Rolando *et al*., 2016). 48 hours after *Drosha* cKO, we administered doxycycline to induce GFP expression from the Tet-on ctrl and Tet-on 3’ UTR HP reporter constructs, and compared GFP expression from these reporters in wild type (Tom^−^) and *Drosha* cKO (Tom^+^) NSCs by FACS analysis (**Figure 2E**). *Drosha* cKO increased GFP expression from the Tet-on 3’ UTR HP reporter compared to cells with intact Drosha alleles. Conversely, *Drosha* cKO did not affect GFP expression from the Tet-on ctrl reporter (**Figure 2G,H**). Therefore, insertion of the NFIB 3’ UTR HP into the Tet-on 3’ UTR HP reporter construct conveyed Drosha sensitivity, and the system enables quantification of Drosha cleavage of target mRNAs.

### Drosha and NFIB RNA interacting proteins are novel regulators of NFIB HP processing

We hypothesized that the Drosha and NFIB mRNA interacting proteins modulate the activity of Drosha towards its targets. To assess these effects, we selected RBPs from the Drosha interacting protein dataset and performed gain-of-function analysis in the Tet-on DG NSCs (**Figure 3A**). We expressed Bub3, Ddx5, Ddx17, DGCR8, Dhx9, Fus, Hnrnpa1, Hnrnpu, Khsrp, Pabpn1, Prpf6, Qki, Safb1, Sam68, Sart1, Sf1, Tdp4 and Trim9 in DG NSCs (*CAG::RBP-IRES-CFP*) or expressed CFP as a control (*CAG::IRES-CFP*), and assessed the regulation of the Tet-on ctrl and Tet-on 3’ UTR HP reporters. 24 hours after expression of the RBP or CFP control, we induced the Tet-on ctrl and Tet-on 3’ UTR HP reporters by doxycycline induction, and quantified GFP expression at the single cell level by FACS and population level after sorting by RT-qPCR (**Figure 3A,B**).

In order to identify RBPs that acted directly on the NFIB 3’ UTR HP sequences and not through indirect effects on the Tet-on EGFPd2 expression cassettes via transcriptional and translational regulation, independent of the NFIB 3’ UTR HP sequences, the changes in GFP intensity following RBP expression in the Tet-on 3’ UTR HP DG NSC line were standardized to changes induced by the same RBP expressed in the Tet-on ctrl DG NSCs. Among the eighteen RBPs tested, Dhx9, Sf1, Hnrnpu and TDP-43 caused the most dramatic increases in GFP expression in a NFIB 3’ UTR HP-dependent fashion. Conversely, Safb1, Qki, Sam68 and Sart1 induced the most pronounced reduction in GFP protein expression (**Figure 3C,D**). Of the 8 RBPs that showed the greatest effects on GFP levels from the Tet-on 3’ UTR HP reporter, Fus and Safb1 were those that also showed a corresponding change in the GFP mRNA levels (Fus 824%, Safb1 48%) suggesting their actions were based on regulation of mRNA turnover (**Figure 3D**).

Drosha destabilizes NFIB mRNA and thus reduces NFIB protein expression and similarly reduces GFP expression from the Tet-on 3’ UTR HP reporter. Therefore, we hypothesized that Drosha partners that reduce GFP mRNA and protein expression from the Tet-on 3’ UTR HP construct are potential positive regulators of Drosha non-canonical activity on NFIB mRNA. Safb1 strongly reduced GFP mRNA and protein expression suggesting its direct involvement in the regulation NFIB mRNA stability through the 3’ UTR HP (**Figure 3D**).

### Safb1 regulates NFIB 3’ UTR HP stability via Drosha

Safb1 showed a strong binding preference towards the NFIB 3’ UTR HP compared to the NFIB 5’ UTR HP (**Figure 1-figure supplement 3E**). We addressed whether this specificity underlies differences in a functional regulation of the NFIB 5’ UTR and 3’ UTR HPs by Drosha. Therefore, we generated an independent stable DG NSC line expressing a Tet-on 5’ UTR HP construct, where the NFIB 3’ UTR HP sequence in the Tet-on 3’ UTR HP construct had been exchanged with the NFIB 5’ UTR HP sequence (**Figure 4A,B** and **Figure 4-figure supplement 1A,B**).

**Figure 4:**
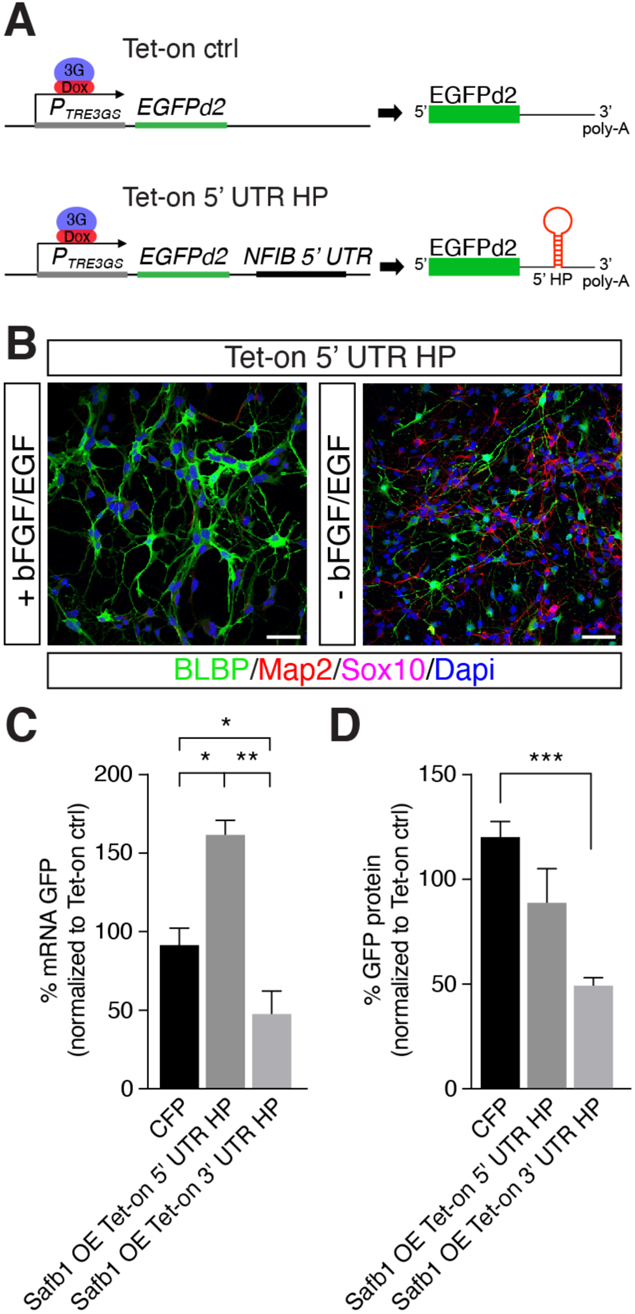
Safb1 overexpression modulates NFIB reporter expression. **A.** Scheme of the constructs used for generating Tet-on ctrl and Tet-on 5’ UTR HP DG NSCs. **B.** Characterization of the Tet-on ctrl and NFIB 5’ UTR HP expressing DG NSCs under expansion (+bFGF/EGF) and differentiation (-bFGF/EGF) culture conditions. Immunohistochemistry for the progenitor marker BLBP, neuronal protein Map2, and oligodendrocyte protein Sox10. Scale bar 50 µm. **C.** RT-qPCR analysis of EGFPd2 (GFP) mRNA levels of CFP expressing and Safb1 overexpressing (Safb1 OE) Tet-on 3’ UTR HP and Tet-on 5’ UTR HP DG NSCs (x-axis). Percent mean EGFPd2 (GFP) mRNA expression by Tet-on 3’ UTR HP and Tet-on 5’ UTR HP DG NSCs normalized to expression by Tet-on ctrl DG NSCs (y-axis). n = 3; one-way ANOVA with Holm-Sidak’s test: *p<0.05, **p<0.01, **p<0.001. Error bars SEM. **D.** FACS analysis of EGFPd2 (GFP) protein fluorescence of CFP expressing and Safb1 overexpressing (Safb1 OE) Tet-on 3’ UTR HP and Tet-on 5’ UTR HP DG NSCs (x-axis). Percent median EGFPd2 (GFP) protein fluorescence intensity of Tet-on 3’ UTR HP and Tet-on 5’ UTR HP DG NSCs normalized to fluorescence intensity of Tet-on ctrl DG (y-axis). n = 6; one-way ANOVA with Tukey’s test: ***p<0.001. Error bars SEM.

**Figure 4-figure supplement 1:**
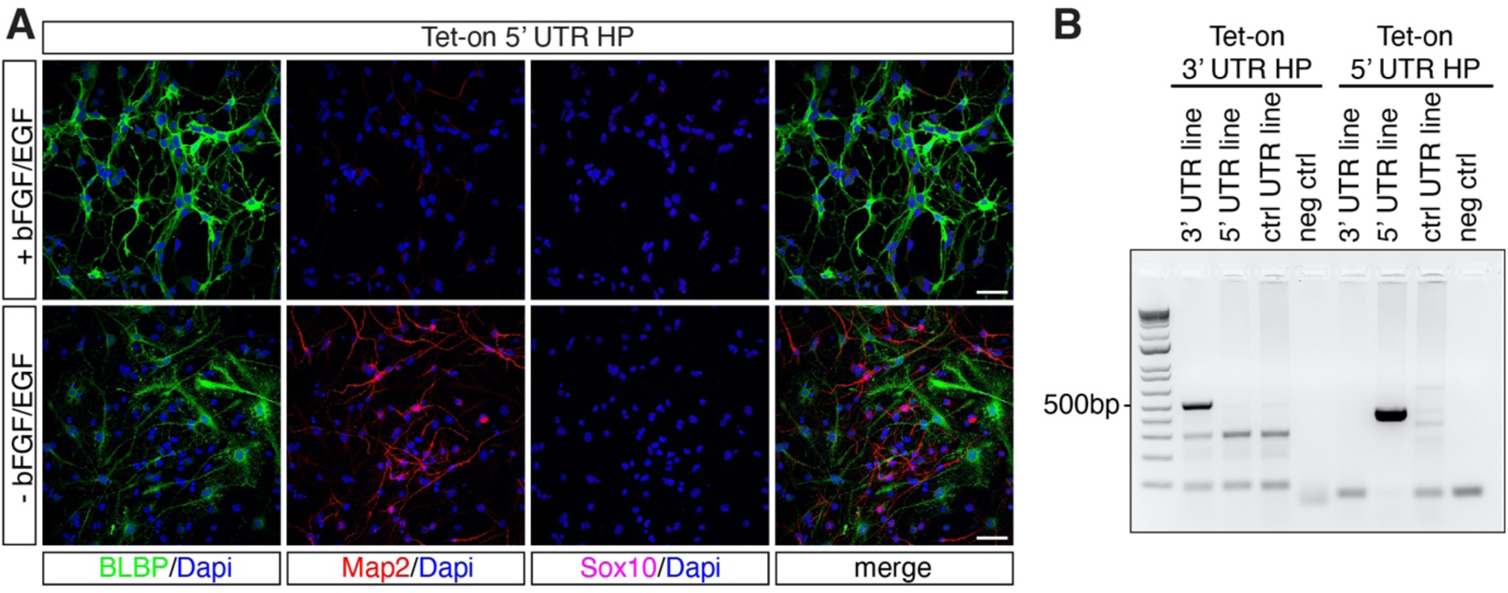
DG NSC NFIB 5’ UTR reporter line retains NSC properties. **A.** DG NSCs under expansion (+bFGF/EGF) and differentiation (-bFGF/EGF) culture conditions. Immunohistochemistry for progenitor marker BLBP, neuronal marker Map2 and oligodendrocyte marker Sox10. Scale bar 50 µm. **B.** Genotyping of the stable Tet-on 3’ UTR HP DG NSCs and Tet-on 5’ UTR HP DG NSCs. Specific amplicons for the Tet-on 3’ UTR HP construct and (514 bp) and 5’ UTR HP construct (430 bp) are found only in the respective lines. left: amplification with primers specific for the NFIB 3’ UTR HP construct, right: amplification with primers specific for the NFIB 5’ UTR HP construct. Negative control: (neg ctrl) H_2_O.

We expressed Safb1 (*CAG::Safb1-IRES-CFP)* in Tet-on ctrl, Tet-on 5’ UTR HP and Tet-on 3’ UTR HP DG NSCs and compared the levels of GFP protein and mRNA 48 hours after doxycycline induction relative to the effects of expressing CFP alone (**Figure 3A**). Safb1 overexpression significantly reduced GFP mRNA expression in Tet-on 3’ UTR HP DG NSCs (**Figure 4C**). Interestingly and in stark contrast, Safb1 overexpression increased GFP mRNA in Tet-on 5’ UTR HP DG NSCs indicating key differential roles of Safb1 on the two HPs (**Figure 4C**). Similarly, GFP protein intensity was also reduced in the Tet-on 3’ UTR HP DG NSCs following Safb1 expression (**Figure 4D**). These data show that Safb1 can regulate NFIB mRNA stability through the 3’ UTR HP.

To address whether Safb1 and Drosha cooperate to regulate endogenous NFIB mRNA, we first investigated whether Drosha and Safb1 physically interact in DG NSCs via a co-IP assay. We immunoprecipitated endogenous Drosha from DG NSCs with an anti-Drosha antibody and detected endogenous Safb1 in the IP but not PKC-*α*, which we used as a negative control as it has not been reported to interact with Drosha and was not identified in the MS of the Drosha IP (**Figure 5A** and **Figure 5-figure supplement 1A-C**). The binding of Safb1 to Drosha was confirmed by Safb1 co-IP (**Figure 5-figure supplement 1D-F**). We assessed whether Dorsha-Safb1 interaction required RNA. Performing the co-IP in the presence of RNAse did not affect Drosha interaction with Safb1 (**Figure 5-figure supplement 1D-F**).

**Figure 5:**
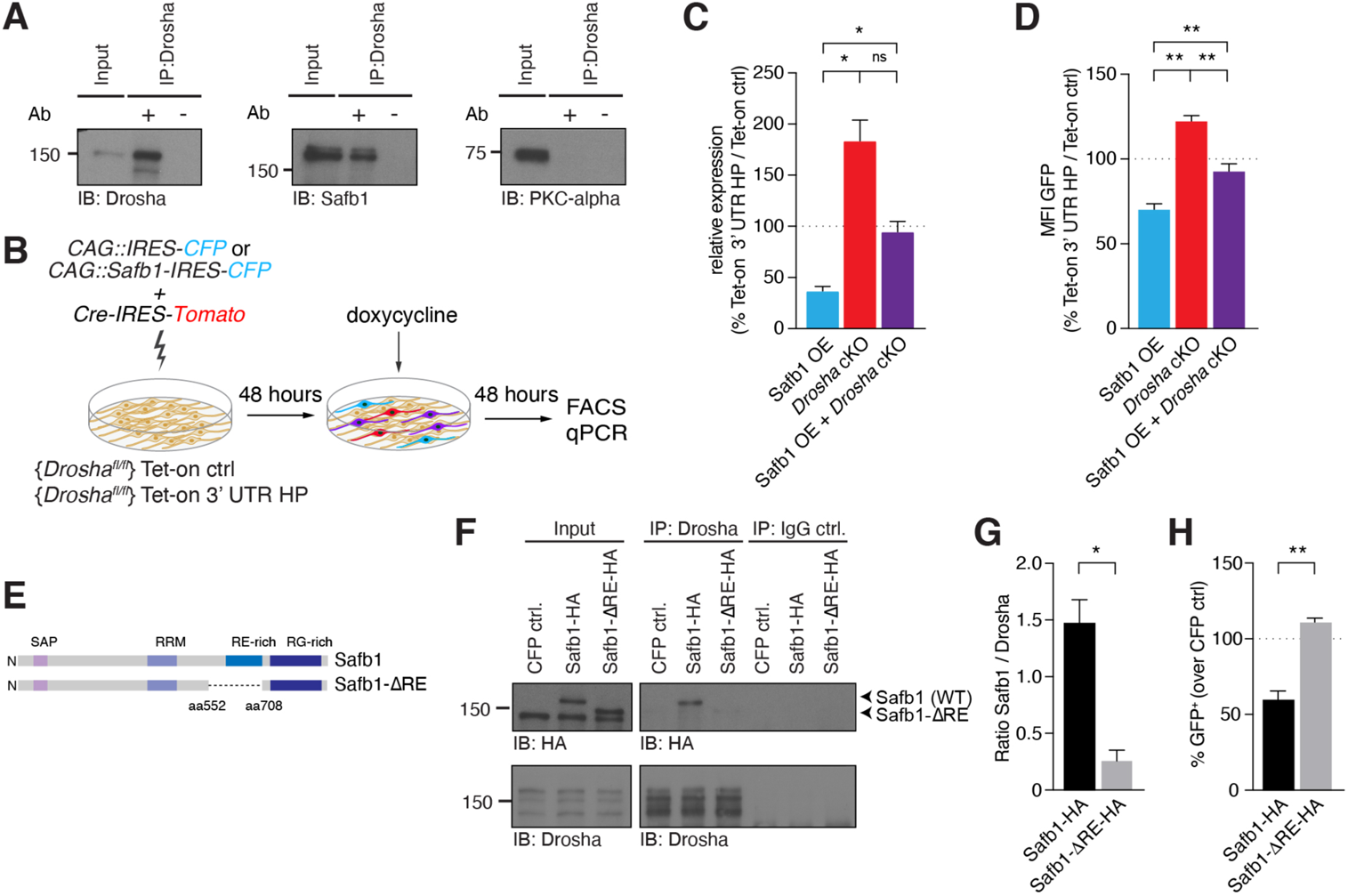
Safb1 mediated reduction in NFIB expression depends on Drosha activity. **A.** Immunoblot validation of Drosha-Safb1 interaction. Drosha IP shows enrichment of Drosha (13% of input) and co-IP of Safb1. Input: 2.5% of total lysate. PKC-*α* was used as negative control for the co-IP. **B.** Scheme of the experimental setup for analysis of the effects of Safb1 overexpression (*CAG::Safb1-IRES-CFP)* on Tet-on ctrl and Tet-on 3’ UTR HP expression in floxed *Drosha* (*Drosha^fl/fl^*) DG NSCs by FACS and RT-qPCR analysis. **C.** RT-qPCR analysis of EGFPd2 (GFP) mRNA levels in Tet-on 3’ UTR HP DG NSCs. Quantification of mean EGFPd2 (GFP) mRNA expression in Tet-on 3’ UTR HP DG NSCs after *Drosha* cKO and overexpression (OE) of Safb1 with/without *Drosha* cKO relative to Tet-on ctrl DG NSCs. 100% line indicates no difference in expression. n = 3, one-way ANOVA with Dunnett’s test: *p<0.05. Error bars SEM. **D.** FACS analysis of EGFPd2 (GFP) protein fluorescence in Tet-on 3’ UTR HP DG NSCs. Quantification of median EGFPd2 intensity (MFI GFP) of Tet-on 3’ UTR HP DG NSCs after *Drosha* cKO and overexpression (OE) of Safb1 with/without *Drosha* cKO relative to Tet-on ctrl DG NSCs. 100% line indicates no difference in expression. n = 5, one-way ANOVA with Tukey’s test: **p<0.01. Error bars SEM. **E**. Schematic representation of the Safb1 and Safb1-ΔRE mutant lacking amino acids 552-708 encoding the RE domain. **F**. Immunoblot (IB) analysis of Safb1-HA (WT), Safb1-ΔRE-HA and Drosha in Drosha-IP and IgG isotype control IP from transfected N2A cells. Input: 2.5% of total lysate. **F.** Quantification of binding capacity of Drosha to Safb1 and Safb1-ΔRE. Binding was quantified as the levels of Safb1-HA or Safb1-ΔRE-HA relative to the Drosha in the respective co-IP measured by densitometry of immunoblots. n = 3, two-tailed Welch’s t-test: *p<0.05. Error bars SEM. **H**. FACS analysis of EGFPd2 (GFP) protein fluorescence in Tet-on 3’ UTR HP DG NSCs following Safb1 and Safb1-ΔRE overexpression. Percentage of median EGFPd2 (GFP) intensity over *CAG::IRES-CFP* control transfection. 100% line indicates no difference in expression. n = 4, two-tailed Mann-Whitney test: **p<0.01. Error bars SEM.

**Figure 5-figure supplement 1:**
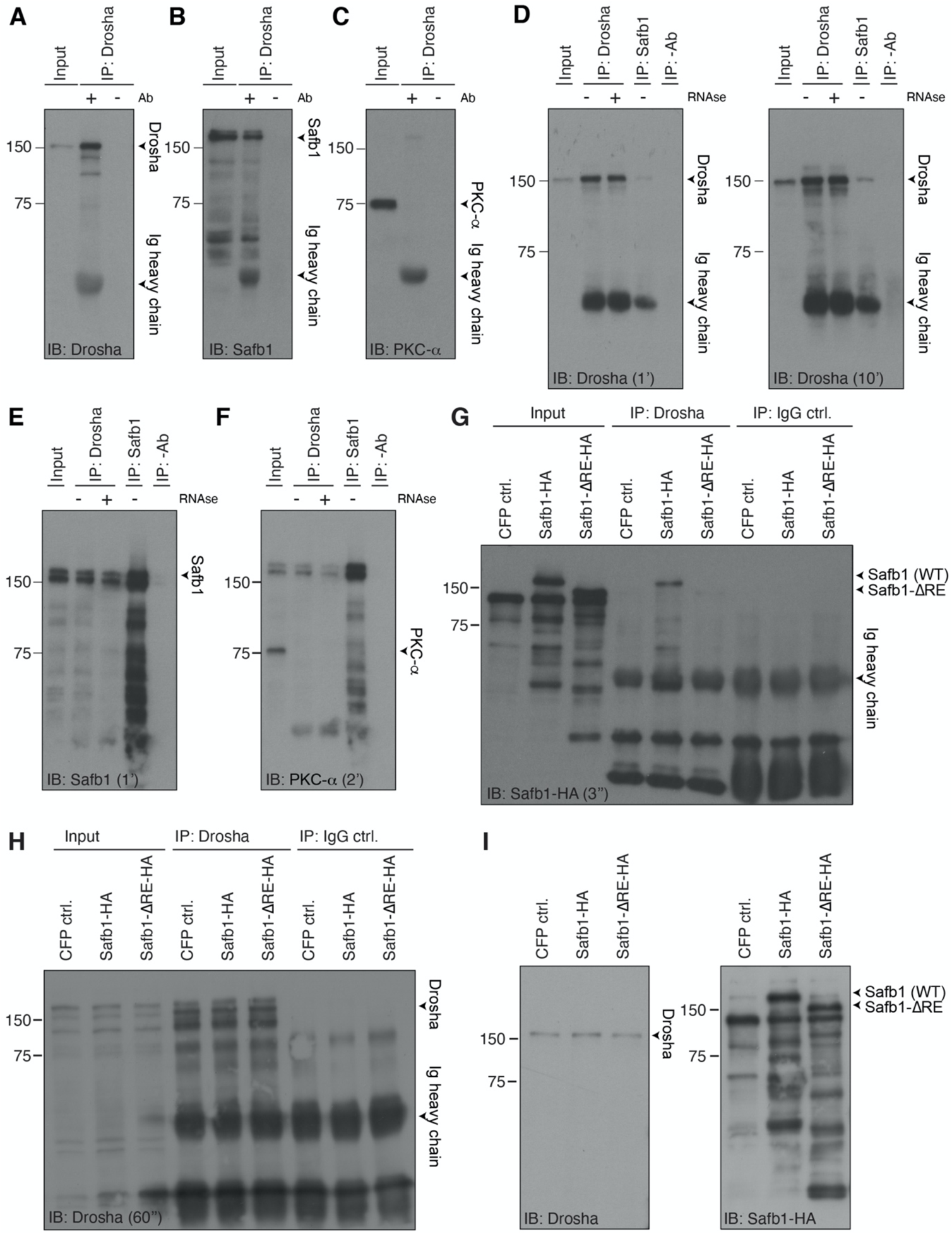
Safb1 and Drosha are able to form a protein complex. **A.** Immunoblot (IB) of Drosha-immunoprecipitation (IP) (related to Figure 5A), showing precipitation of Drosha. Immunoglobulin (Ig) heavy chain of the precipitating antibody detected in the immunoblots. Input: 2.5% of the lysate used in the IP. Negative control precipitation without antibody (-Ab). **B**. Immunoblot (IB) of Drosha-IP (related to Figure 5A), showing coprecipitation of Safb1. Immunoglobulin (Ig) heavy chain of the precipitating antibody detected in the immunoblots. Input: 2.5%. Negative control precipitation without antibody (-Ab). **C**. Immunoblot (IB) of Drosha-IP (related to Figure 5A), showing no precipitation of PKC-*α*. Immunoglobulin (Ig) heavy chain of the precipitating antibody detected in the immunoblots. Input: 2.5%. Negative control precipitation without antibody (-Ab). **D**. Immunoblot (IB) for Drosha of Drosha-IP performed in the presences (+) or absence (-) of RNAse and Safb1-IP. Immunoglobulin (Ig) heavy chain of the precipitating antibody detected in the immunoblots. Input: 2.5%. Negative control precipitation without antibody (-Ab). Exposure times - 1 minute and 10 minutes. **E**. Immunoblot (IB) for Safb1 of Drosha-IP and Safb1-IP performed in the presences (+) or absence (-) of RNAse. Input: 2.5% of the lysate used in the IP. Negative control precipitation without antibody (-Ab). Exposure time - 1 minute. **F**. Immunoblot (IB) for PKC-*α* of Drosha-IP and Safb1-IP performed in the presences (+) or absence (-) of RNAse, showing no precipitation of PKC-*α* (note: re-blotted after Safb1 detection without stripping of the blot). Input: 2.5% of the lysate used in the IP. Negative control precipitation without antibody (-Ab). Exposure time - 2 minutes. **G.** Immunoblot (IB) with anti-HA antibodies of Drosha-IP and control IgG-IP of transfected N2A cells expressing CFP ctrl. (not HA-tagged), Safb1-HA or Safb1-ΔRE-HA. Input: 2.5% of the lysate used in the IP. Negative control precipitation without with IgG isotype control antibody (IgG ctrl.). **H.** Immunoblot (IB) for Drosha of Drosha-IP and control IgG-IP of transfected N2A cells expressing CFP ctrl. (not HA-tagged), Safb1-HA or Safb1-ΔRE-HA showing precipitation of endogenous Drosha. Analysis confirms equal levels of Drosha in all three samples in input and IP, respectively. Input: 2.5% of the lysate used in the IP. Negative control precipitation without with IgG isotype control antibody (IgG ctrl.). **I.** Immunoblot (IB) for endogenous Drosha as loading control and with anti-HA antibodies of transfected DG NSCs cells expressing CFP ctrl. (not HA-tagged), Safb1-HA (migrating at ∼kDa 160) or Safb1-ΔRE-HA (migrating at ∼kDa 145).

To address whether Safb1 and Drosha cooperate at the NFIB 3’ UTR HP, we analyzed the effects of Safb1 overexpression on Tet-on 3’ UTR HP reporter activity in the presence and absence of Drosha (**Figure 5B**). In the presence of Drosha, Safb1 overexpression reduced GFP expression from the doxycycline-induced Tet-on 3’ UTR HP reporter at the RNA (RT-qPCR) and protein levels (FACS), compared to the Tet-on GFP reporter (**Figure 5B-D**). Thus, Safb1 negatively regulates the expression of a NFIB 3’ UTR HP containing mRNA. We then addressed whether the effects of Safb1 overexpression depend on Drosha activity by expressing Safb1 combined with *Drosha* cKO. While *Drosha* cKO increased GFP expression from the Tet-on 3’ UTR HP reporter at the mRNA and protein levels, simultaneous *Drosha* cKO and overexpression of Safb1 reversed the effects on both mRNA and protein expression (**Figure 5B-D**). Therefore, Drosha and Safb1 work together to regulate Tet-on 3’ UTR HP expression.

We addressed whether Drosha processing of NFIB 3’ UTR HP is dependent upon binding to Safb1. The RE-rich region of Safb2 has been shown to bind Drosha (Hutter et al., 2020). Therefore and in analogy, we generated a Safb1 mutant lacking 156 amino acids containing the RE-rich region of Safb1 (aa552 to aa708) (**Figure 5E**). To assess Drosha binding of the Safb1-ΔRE protein, we tagged the mutant and wild type Safb1 variants with HA tags to distinguish them from endogenous protein, expressed these in N2A cells, and performed a Drosha co-IP followed by anti-HA immunoblot. We quantified the amount of Safb1 wild type or Safb1-ΔRE mutant coprecipitated with Drosha (**Figure 5F** and **Figure 5-figure supplement 1G,H**). Safb1-ΔRE showed a 5.7-fold reduction in binding to Drosha compared to Safb1 wild type protein (**Figure 5G**). We addressed the functional consequences of deleting the RE domain of Safb1 on NFIB 3’ UTR regulation and expressed the Safb1-ΔRE mutant in Tet-on 3’ UTR HP and Tet-on ctrl DG NSCs. Strikingly, and unlike overexpression of the wild type HA-tagged Safb1 protein, overexpression of the Safb1-ΔRE mutant did not reduce the expression of GFP in Tet-on 3’ UTR HP compared to Tet-on ctrl DG NSCs (**Figure 5H**), even though the Safb1-ΔRE protein was expressed at similar levels to the wild type Safb1 (**Figure 5-figure supplement 1I**). These data indicated that the Safb1 RE domain is involved in Drosha binding and enhances Drosha activity on the NFIB 3’ UTR HP.

### Safb1 represses NFIB expression in DG NSCs

Safb1 reduced expression of the NFIB 3’ UTR HP reporter in a Drosha-dependent manner. We mapped Safb1 RNA binding motifs and found several potential binding sites in the NFIB 3’ UTR HP and flanking sequence (**Figure 6A**) (Rivers *et al*., 2015; Van Nostrand et al., 2020). NFIB mRNA was recently identified as a SAFB1 target in HepG cells and the interaction domain mapped to the 3’ UTR region of NFIB mRNA overlapping the 3’ UTR HP (**Figure 6A** and **Figure 6-figure supplement 1A**) (Van Nostrand *et al*., 2020). We evaluated Safb1 binding to the endogenous NFIB mRNA by crosslinking and immunoprecipitation (CLIP) with anti-Safb1 antibodies from DG NSCs followed by RT-qPCR analysis (**Figure 6B**). Both NFIB mRNA and a known Safb1 target, Hnrnpu mRNA, were bound and precipitated with Safb1 (**Figure 6C**). Therefore, we evaluated the effects of Safb1 overexpression on the levels of endogenous NFIB mRNA in DG NSCs by RT-qPCR. Safb1 overexpression reduced NFIB mRNA levels (**Figure 6D**) (Knuckles *et al*., 2012; Rolando *et al*., 2016). Thus, Safb1 directly binds NFIB mRNA and represses its levels in DG NSCs.

**Figure 6:**
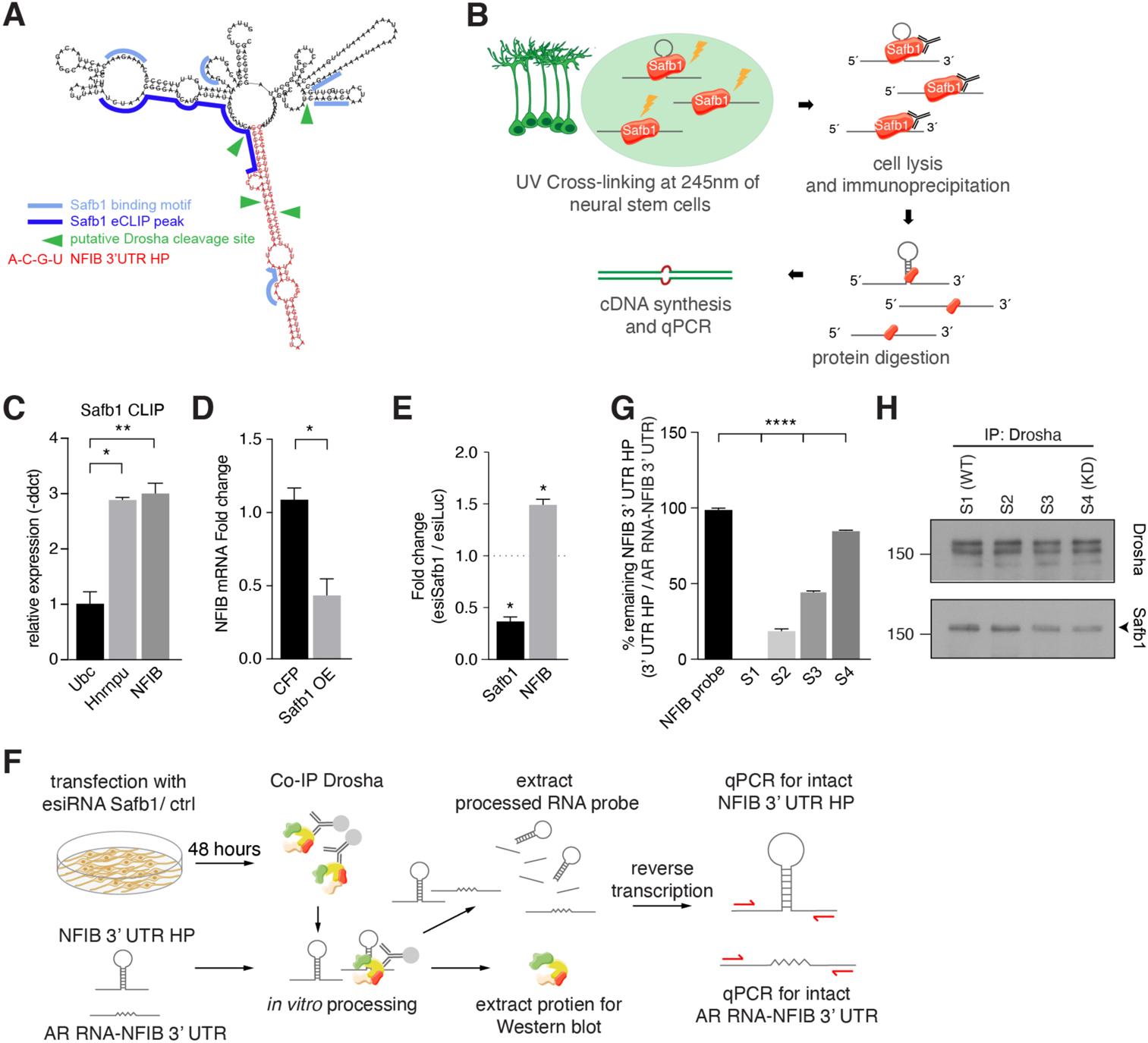
Safb1 binds and regulates endogenous NFIB expression. **A.** *In silico* motif analysis of Safb1 binding sites (light and dark blue) and the predicted secondary structure of the NFIB 3’ UTR HP (red). SAFB1 eCLIP peak mapped on the NFIB mRNA in HepG cells by Van Nostrand et al. (Van Nostrand *et al*., 2020). Putative Drosha cleavage sites (green arrowheads) mapped in Rolando et al. 2016. **B.** Scheme of the experimental setup for Safb1 cross-linked immunoprecipitation (CLIP) from DG NSCs and detection of bound RNA transcripts by RT-qPCR. **C.** RT-qPCR analysis of Safb1 CLIP from DG NSCs. Relative levels of Ubc, Hnrnpu and NFIB pull-down (-ddct values) calculated over input and minus antibody (-Ab) control. Negative control: Ubc, Positive control: Hnrnpu. n = 3, one-way ANOVA with Dunnett’s test: *p<0.05, **p<0.01. Error bars SEM. **D.** RT-qPCR analysis of NFIB mRNA levels in DG NSCs transfected with CFP or Safb1 overexpression (OE) vectors displayed as fold change NFIB mRNA over untransfected cells. n = 4, two-tailed Mann-Whitney test: *p<0.05. Error bars SEM. **E.** Quantification of relative Safb1 and NFIB mRNA expression levels in Safb1 knockdown (esiSafb1) N2A cells after 48 hours compared to control (esiLuc) transfected cells. Two-tailed Students t-test: *p<0.05. Error bars SEM. **F.** Scheme of the experimental setup for Drosha complex immunoprecipitation from N2A cells with/without Safb1 knockdown (esiSafb1) and *in vitro* processing of NFIB 3’ UTR HP and AR RNA-NFIB 3’ UTR (control) RNA probes. Drosha complexes were immunoprecipitated with anti-Drosha antibodies under native conditions from Safb1 esiRNA knockdown (esiRNA Safb1) N2A and untransfected (control) N2A cells. Drosha complexes from control and Safb1 knockdown cells were mixed (ratios 100:0, 66:33, 33:66 and 0:100) and incubated with 500 ng NFIB 3’ UTR HP RNA or AR RNA-NFIB 3’ UTR RNA probe for 30 minutes. Proteins and RNA were extracted and analyzed by immunoblot and RT-qPCR for intact NFIB 3’ UTR HP RNA and AR RNA-NFIB 3’ UTR RNA. Both the NFIB 3’ UTR HP RNA and AR RNA-NFIB 3’ UTR RNA probe share the same primer binding sequences. Although the *in vitro* processing experiment was performed in the presence of RNAse inhibitor, the AR RNA-NFIB 3’ UTR RNA probe served as a control for non-specific RNAse activity. **G.** RT-qPCR analysis of intact NFIB 3’ UTR HP RNA probe after incubation with Drosha complexes (S1-S4). Unprocessed probe NFIB 3’ UTR HP (NFIB probe) was used to calculate the percent remaining intact probe in samples S1-S4. The levels of intact NFIB 3’ UTR HP probe were normalized to the levels of AR RNA-NFIB 3’ UTR hybrid probe (AR RNA-NFIB 3’ UTR) incubated with the same lysates and compared to input unprocessed NFIB probe levels (see Figure 6-figure supplement 1F,G). n = 3, one-way ANOVA with Tukey’s test: ****p<0.0001. Error bars SEM. **H**. Immunoblot analysis of Drosha and Safb1 in the Drosha-IP complexes in samples S1-S4 of the *in vitro* NFIB 3’ UTR HP and AR RNA-NFIB 3’ UTR probe processing experiments. Sample S1 has endogenous levels of Safb1, S4 is a Safb1 KD sample. S2 and S3 are mixes of S1 and S4 in ratios 66:33 and 33:66, respectively. Drosha levels were constant in S1-S4.

**Figure 6-figure supplement 1:**
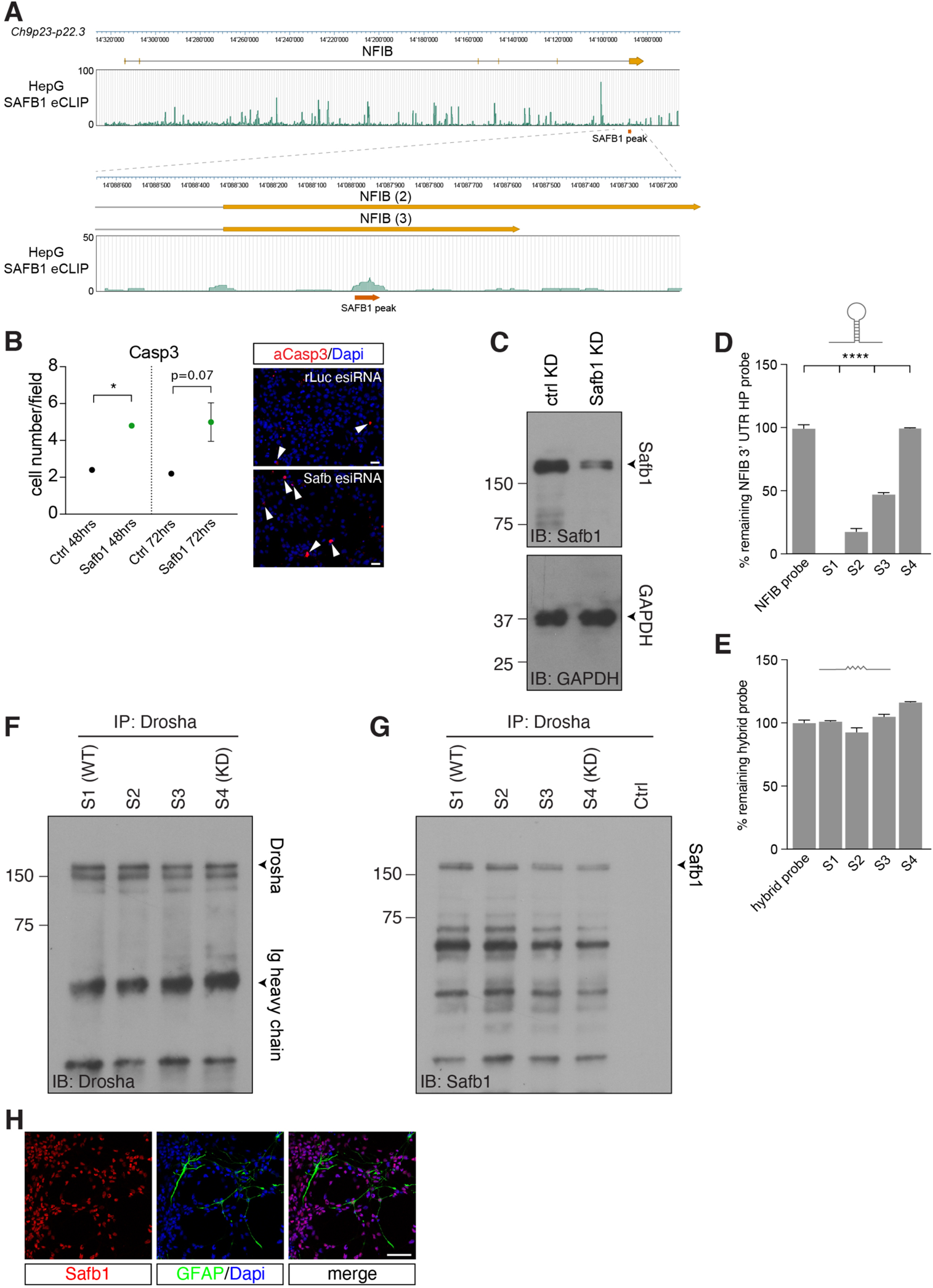
Safb1 contributes to regulation of NFIB mRNA level. **A.** Mapping of SAFB1 eCLIP peaks from HepG cells to the *NFIB* in the minus strand of chromosome 9p23-p22.3 by Van Nostrand et al. (Van Nostrand *et al*., 2020). Lower panel shows the peak call for SAFB1 binding in the two alternative 3’ UTR regions of NFIB (transcript 2 and 3), which corresponds precisely to the hairpin region and the CLIP site shown by Rivers et al. (Figure 6A) (Rivers *et al*., 2015). **B.** Quantification of activated Caspase 3 expression in Safb1 esiRNA knockdown and control (esiRNA rLuc) transfected cells after 48 hours, and 72 hours. Safb1 knockdown results in a rapid induction of apoptosis. Two-tailed Students t-test: *p<0.05. Error bars SEM. **C**. Immunoblot (IB) for Safb1 in ctrl KD (esiRNA rLuc) and Safb1 KD (esiRNA Safb1) lysates (related to Figure 6F). Densitometric analysis revealed 70% reduction in Safb1 protein levels in the Safb1 knockdown (Safb1 KD) sample compared to control (ctrl: esiRNA rLuc). Loading control Immunoblot (IB) for GAPDH. **D.** RT-qPCR analysis of intact NFIB 3’ UTR HP RNA probe after incubation with Drosha-IP complexes. Unprocessed probe NFIB 3’ UTR HP (NFIB probe) was used to calculate the percent remaining of intact probe in samples S1-S4 (control and Safb1 knockdown Drosha complexes mixed at ratios S1-100:0, S2-66:33, S3-33:66 and S4-0:100, respectively). *In vitro* processing reactions were performed in the presence of RNAse inhibitor. n = 3, one-way ANOVA with Tukey’s test: ****p<0.0001. Error bars SEM. **E.** RT-qPCR analysis of intact AR RNA-NFIB 3’ UTR hybrid probe after incubation with Drosha-IP complexes. Unprocessed AR RNA-NFIB 3’ UTR hybrid probe (hybrid probe) was used to calculate the percent remaining, intact probe in samples S1-S4 (control and Safb1 knockdown Drosha complexes mixed at ratios S1-100:0, S2-66:33, S3-33:66 and S4-0:100, respectively). *In vitro* processing reactions were performed in the presence of RNAse inhibitor. **F**. Immunoblot (IB) analysis of Drosha in Drosha-IP complexes in samples S1-S4. Sample S1 is wild type Safb1, S4 corresponds to Safb1 KD sample. S2 and S3 are mixed S1 and S4 in ratios 66:33 and 33:66, respectively. Drosha levels were similar in S1-S4 used in *in vitro* NFIB 3’ UTR HP and hybrid NFIB-AR probe processing experiments. **G.** Immunoblot (IB) analysis of Safb1 expression in S1-S4 confirming a linear reduction in Safb1 levels across the samples. Crtl - no Drosha-IP complex. **H.** Safb1 expression in cultured DG NSCs. Immunofluorescent staining for Safb1 and GFAP. Scale bar 50 µm.

As Safb1 promotes Drosha cleavage of NFIB transcripts and Drosha activity controls DG NSC fate *in vivo* and *in vitro* (Rolando *et al*., 2016), we evaluated the role of Safb1 in DG NSCs by esiRNA-mediated knockdown (KD) *in vitro*. Safb1 KD in DG NSCs led to a rapid increase in activated Casp3 and cell death within 48 hours, preventing further fate analysis (**Figure 6-figure supplement 1B**). Therefore, we turned to neuroblastoma cells (N2A) that express Drosha, Safb1 and NFIB mRNA. Safb1 KD caused a drastic increase in NFIB mRNA levels in N2A cells underlining the importance of Safb1 for efficient NFIB mRNA processing (**Figure 6E** and **Figure 6-figure supplement 1C**).

We assessed whether Safb1 directly affects NFIB 3’ UTR HP cleavage by performing a modified *in vitro* processing assay (Rolando *et al*., 2016). *In vitro* transcribed NFIB 3’ UTR HP RNA probe was incubated with the Drosha/Safb1 complexes precipitated from N2A cells and processing via Drosha assessed by RT-qPCR to detect intact probe (**Figure 6F,G**). We also used the control AR RNA-NFIB 3’ UTR probe, which contains the regions flanking the NFIB 3’ UTR HP but where the HP is replaced by the AR sequence, which is not cleaved by Drosha, in the *in vitro* processing experiments to monitor potential non-specific RNAse activity. The NFIB 3’ UTR HP and AR RNA-NFIB 3’ UTR probes share the same PCR primer sequences allowing the same PCR conditions to be used to quantify both probes. The NFIB 3’ UTR HP probe was completely processed following incubation with Drosha complexes (S1 (WT) in **Figure 6F** and **Figure 6-figure supplement 1D**). However, the AR RNA-NFIB 3’ UTR control probe remained intact (S1 (WT) in **Figure 6-figure supplement 1E**).

To address whether the levels of Safb1 affect Drosha processing of the NFIB 3’ UTR HP, we IPed Drosha complexes from Safb1 knockdown (esiSafb1) N2A cells and mixed these in ratios of 33:66, 66:33 and 100:0 (S2, S3 and S4) with Drosha complexes IPed from N2A cells expressing wild type levels of Safb1. We quantified the levels of Safb1 and Drosha in these reactions by immunoblot and found the expected gradient of Safb1 from S1-S4, and that Drosha levels were constant (**Figure 6H** and **Figure 6-figure supplement 1F,G**). Reducing the levels of Safb1 in the *in vitro* processing reaction reduced the processing of the NFIB 3’ UTR HP probe without affecting AR RNA-NFIB 3’ UTR probe integrity (**Figure 6G,H** and **Figure 6-figure supplement 1D,E**). Drosha complexes from Safb1 knockdown N2A cells did not cleave the NFIB 3’ UTR HP probe which remained intact over the incubation period (S4 (KD) in **Figure 6G,H**). Thus, NFIB 3’ UTR HP processing by Drosha shows a direct relationship to the levels of Safb1 in the complex.

### Safb1 regulates oligodendrocyte differentiation from NSCs

NSCs of the DG generate predominately granule neurons as well as astrocytes but not oligodendrocytes, and their fate restriction is controlled in part by Drosha and its post-transcriptional repression of NFIB expression (Rolando *et al*., 2016). To assess Safb1 expression in the hippocampus, we genetically labelled adult neurogenic DG NSCs by treating *Hes5::CreER^T2^* mice carrying a *Rosa26-CAG::EGFP* Cre-reporter for 5 days with Tamoxifen (Lugert et al., 2012). The Notch signal target *Hes5* is expressed by NSCs in the DG and *Hes5::CreER^T2^* activity can be used to lineage trace neurogenic and gliogenic cells in the adult mouse following Tamoxifen induction (Lugert et al., 2010; Lugert *et al*., 2012). Safb1 protein was expressed by most cells in the adult DG including GFAP^+^, Hes5^+^ radial NSCs (**Figure 7A**). In line with this, DG NSC cultures also showed high levels of Safb1 expression indicating that Safb1 is expressed by DG NSCs both *in vivo* and *in vitro* (**Figure 6-figure supplement 1H**). Safb1 levels were significantly higher in neurons than in astrocytes (GFAP^+^) and oligodendrocytes (Sox10^+^) in the DG and striatum adjacent to the V-SVZ of the lateral ventricle wall, confirming lower levels of Safb1 in glia (**Figure 7A**).

**Figure 7:**
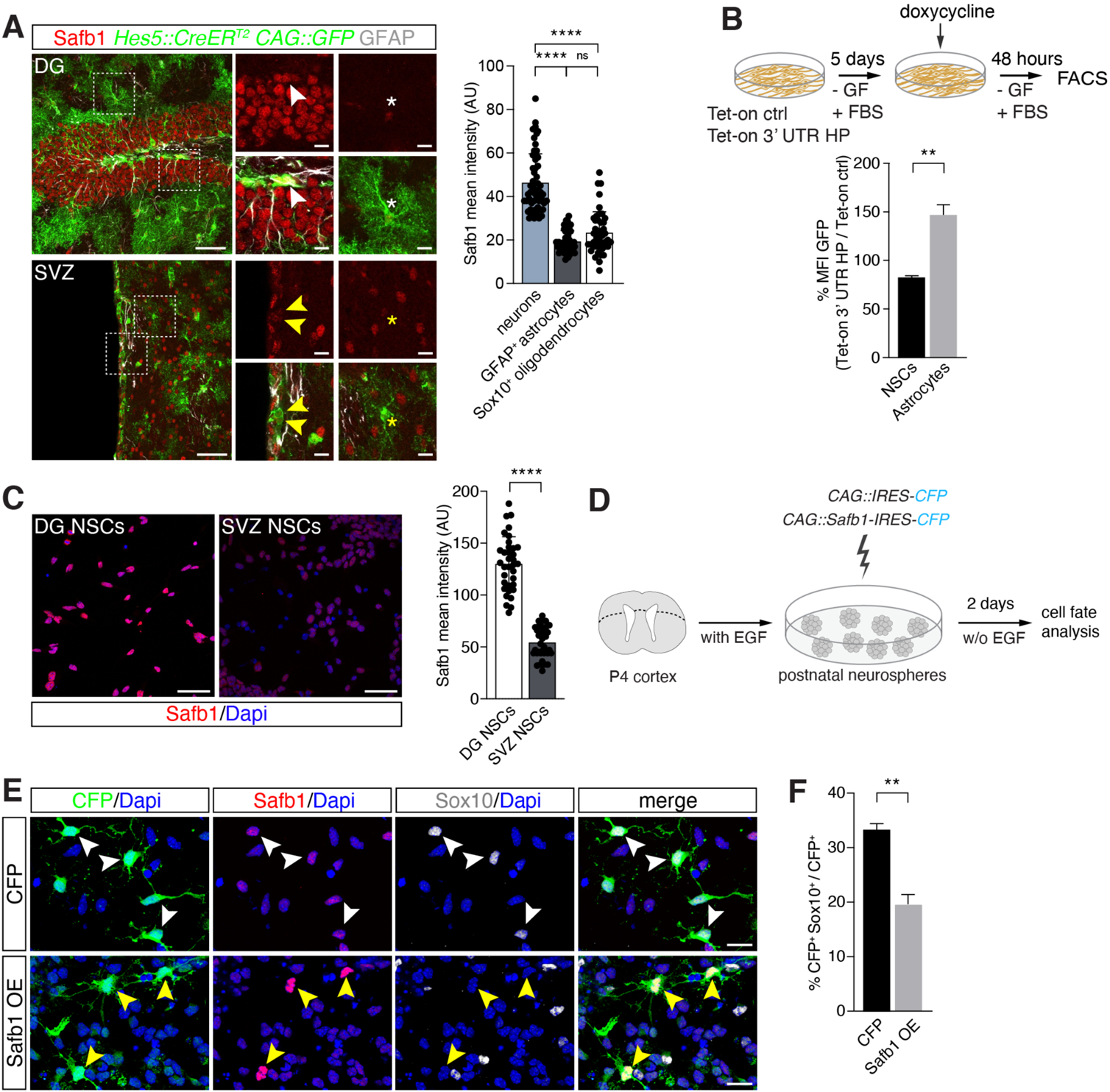
Safb1 overexpression regulates oligodendrogenesis. **A.** Safb1 expression in adult Hes5^+^ DG NSC marked with *Hes5::CreER^T2^* and *CAG::GFP* Cre-reporter (upper panels white arrowheads) and V-SVZ NSCs genetically labelled with *Hes5::CreER^T2^* and *CAG::GFP* Cre-reporter (lower panels yellow arrowheads) *in vivo*. NSCs were labelled *in vivo* by Tamoxifen treatment of *Hes5::CreER^T2^ CAG::GFP* mice (see methods). High magnification images were taken are regions indicated. DG NSCs (GFP^+^; white arrowheads upper panels) and DG neurons (GFP^−^ cells) expresses higher levels of Safb1 than astrocytes (white *). V-SVZ NSCs (GFP^+^; yellow arrowheads lower panels) and astrocytes (yellow *) express lower levels of Safb1 than striatal neurons (GFP^−^ cells). Quantification of Safb1 expression by neurons, astrocytes and oligodendrocytes *in vivo* based on mean intensity levels (arbitrary units). Scale bars: low magnification images 100 µm, high magnification images 20 µm. One-way ANOVA with Tukey’s test: ****p<0.0001, ns - not significant. Error bars SEM. **B.** Scheme of the experimental setup for quantification of EGFPd2 (GFP) protein fluorescence comparing Tet-on 3’ UTR HP over Tet-on ctrl DG NSCs by FACS in undifferentiated NSC state and after growth factor (GF) removal and addition of fetal bovine serum (FBS) for 5 days to induce differentiation to astrocytes followed by a 48-hour doxycycline induction. Quantification of the median EGFPd2 (GFP) intensity of Tet-on 3’ UTR HP relative to Tet-on ctrl in NSCs and astrocytes. n = 3, two-tailed Students t-test: **p<0.01. Error bars SEM. **C.** Images and quantification of Safb1 protein expression by adult DG NSCs and V-SVZ NSCs *in vitro*. Scale bar: 20 µm. Kolmogorov-Smirnov test: ****p<0.0001. Error bars SEM. Scale bars 50µm. **D.** Scheme for the experimental setup for Safb1 (*CAG::Safb1-IRES-CFP*) and control CFP (*CAG::IRES CFP*) overexpression in postnatal V-SVZ NSCs (postnatal day 4: P4) grown as neurospheres in the presence of EGF and cell fate analysis 2 days after EGF withdrawal (w/o EGF) to induces differentiation. **E.** Analysis of the effects of Safb1 overexpression (OE) in V-SVZ neurospheres and effects on oligodendrocyte (Sox10^+^) differentiation compared to CFP (control) expression. Scale bars 20 µm. **F.** Quantification of the percentage of transfected cells (CFP^+^) expressing Sox10^+^ (oligodendrocytes) after Safb1 overexpression (OE) (n = 4) or CFP alone (n = 3). Two-tailed Welch’s t-test: **p<0.01. Error bars SEM.

**Figure 7-figure supplement 1:**
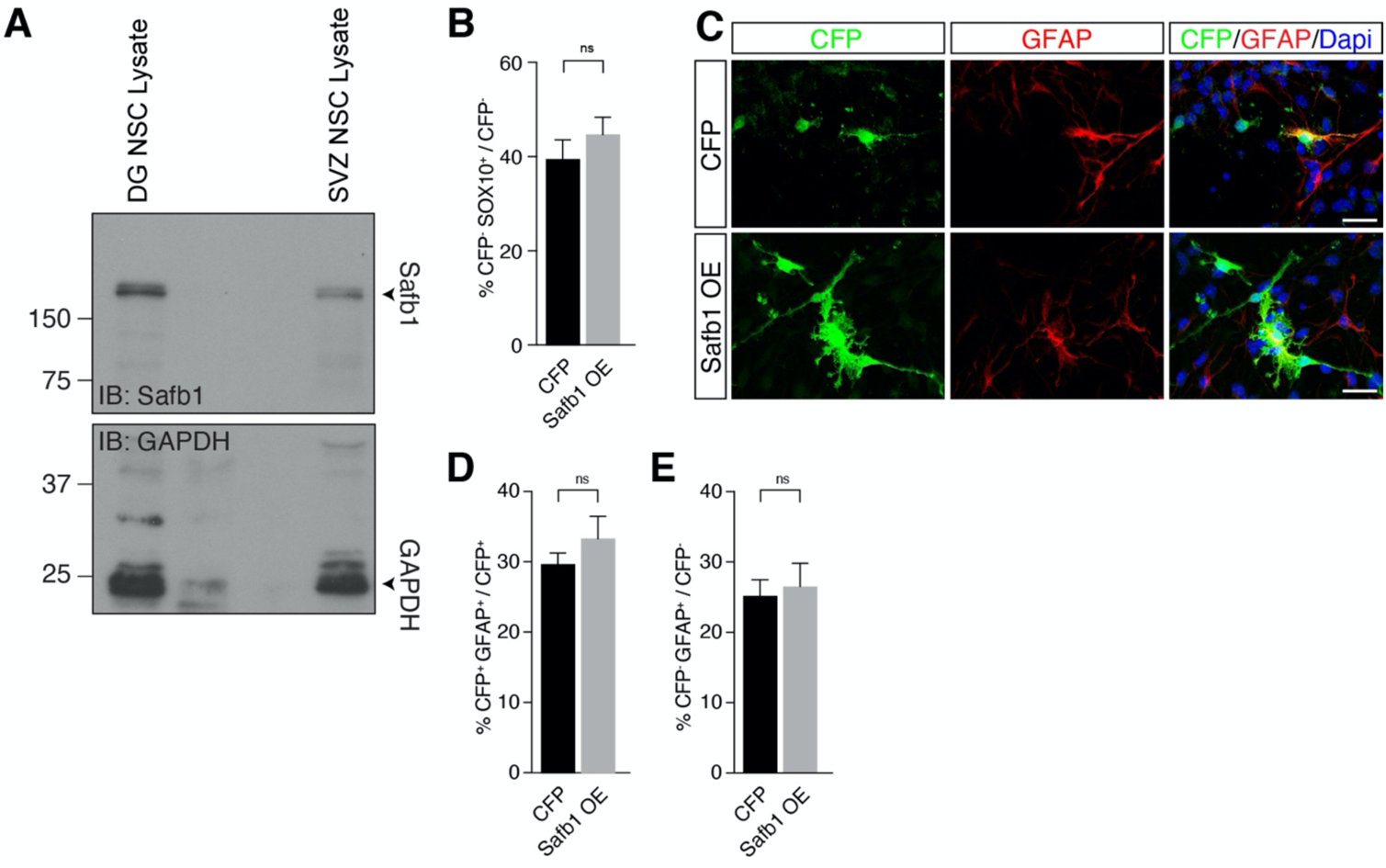
Safb1 overexpression does not increase astrocyte differentiation. **A.** Immunoblot (IB) of cultured DG and ventricular subventricular zone (SVZ) NSCs (related to Figure 7C). Densitometric analysis indicates lower Safb1 expression in the SVZ NSCs (49% of DG NSCs) compared to DG NSCs. Loading control Immunoblot (IB) for GAPDH. **B.** Quantification of immunohistochemistry for CFP negative cells (related to Figure 7E). Mean percentage of CFP^−^Sox10^+^ cells in the CFP control transfected (*CAG::IRES-CFP*, n = 3) and Safb1 OE (*CAG::Safb1-IRES-CFP*, n = 4) samples are shown as percentage of untransfected (CFP^−^). Two-tailed Welch’s t-test: **p<0.01. Error bars SEM. **C**. Safb1 overexpression in perinatal ventricular subventricular zone (SVZ) NSC neurospheres. Cells were transfected with CFP (*CAG::IRES-CFP*) or Safb1 OE vectors (*CAG::Safb1-IRES-CFP*). Staining for CFP and GFAP. Scale bar: 20 µm. **D.** Quantification of immunohistochemistry for CFP positive cells. Mean percentage of CFP^+^GFAP^+^ cells in the CFP control transfected (*CAG::IRES-CFP*, n = 3) and Safb1 OE (*CAG::Safb1-IRES-CFP*, n = 4) samples are shown as percentage of transfected cells (CFP^+^). Two-tailed Welch’s t-test: **p<0.01. Error bars SEM. **E.** Quantification of immunohistochemistry for CFP negative cells. Mean percentage of CFP^−^GFAP^+^ cells in the CFP control transfected (*CAG::IRES-CFP*, n = 3) and Safb1 OE (*CAG::Safb1-IRES-CFP*, n = 4) samples are shown as percent of untransfected (CFP^−^). Two-tailed Welch’s t-test: **p<0.01. Error bars SEM.

As NFIB is associated with driving glial differentiation, we assessed whether the differentiation of NSCs to astrocytes is associated with a decrease in NFIB 3’ UTR HP processing. Therefore, we analyzed the levels of GFP expression from the Tet-on 3’ UTR HP and Tet-on ctrl reporters in NSCs before and after inducing glial differentiation (**Figure 7B**). As predicted, GFP levels from the Tet-on 3’ UTR HP reporter increased in the glial state compared to those from the Tet-on ctrl reporter (**Figure 7B**).

To address the role of Safb1 in NSC fate choices, we performed gain-of-function experiments in V-SVZ NSCs. While adult DG NSCs are fate restricted to generate neurons and astrocytes at the expense of oligodendrocytes, adult V-SVZ NSCs generate all three neural lineages (Kang et al., 2019; Lachapelle et al., 2002; Rolando *et al*., 2016; Sohn et al., 2012). We isolated V-SVZ NSCs from postnatal mice and expanded them *in vitro*. V-SVZ NSCs *in vitro* as *in vivo* expressed Safb1 at lower levels than DG NSCs indicating an inverse correlation between Safb1 expression and oligodendrocytic differentiation potential (**Figure 7A,C** and **Figure 7-figure supplement 1A**).

Overexpression of Safb1 (*CAG::Safb1-IRES-CFP*) in V-SVZ NSCs by nucleofection reduced oligodendrocyte (Sox10^+^CFP^+^) differentiation within 48 hours compared to *CAG::IRES-CFP* expressing control cells (Safb1 overexpression: 19.9 ± 1.6% versus CFP ctrl: 33.2 ± 1.2%: p < 0.01) and non-transfected CFP^−^ cells in the same cultures (**Figure 7D-F** and **Figure 7-figure supplement 1B**). Interestingly, quantification of CFP^+^GFAP^+^ cells revealed no changes in the astrocytic cell population compared to *CAG::IRES-CFP* expressing control cells and non-transfected CFP^−^ cells in the same cultures (**Figure 7-figure supplement 1C-E**). Therefore, Safb1 repressed oligodendrocytic differentiation of multipotent V-SVZ NSCs in a cell autonomous fashion.

## Discussion

A precise control of stem cell fate during tissue development and homeostasis as well as in cancer is critical. Post-transcriptional regulation of gene expression contributes significantly to proteome regulation including in NSCs (Pilaz and Silver, 2015; Ratti et al., 2006). Drosha is the catalytic component of the microRNA Microprocessor where it works together with a dimer of the RBP DGCR8 (Han *et al*., 2004; Nguyen *et al*., 2015). However, more and more Drosha associated proteins are being identified and these form megadalton complexes containing more than 20 different proteins (Macias *et al*., 2015; Nguyen *et al*., 2015). The functions of the larger Drosha complexes remain unclear.

We have identified 165 proteins that interact with Drosha in DG NSCs, revealing an unprecedented complexity in the Drosha interactome. Most of the Drosha-interacting proteins are RBPs, consistent with modulating Drosha activity on its RNA substrates (Gerstberger *et al*., 2014; Huang *et al*., 2018). However, some of the Drosha-associated proteins have not been shown to bind RNA and may be components of larger macromolecular complexes with diverse regulatory functions on Drosha activity. 15% of the Drosha-binding partners in NSCs also associate with Drosha in HEK293T cells indicating some molecular conservation across cell-types (Macias *et al*., 2015; Rouillard *et al*., 2016). However, many of the proteins we identified likely form cell type-specific Drosha complexes. Thus, although the biological functions of most of the Drosha interacting proteins in NSCs are not known, and their roles in controlling Drosha activity remain unclear, our findings provide detailed insights into the complexity and potentially novel functions of Drosha signaling and of its partners.

We previously demonstrated critical, microRNA-independent roles for Drosha in the control of NSC maintenance and in shaping cell fate choices in the developing and adult brain, and others have shown similar functions in other systems (Chong *et al*., 2010; Han et al., 2009; Kim *et al*., 2017; Knuckles *et al*., 2012; Rolando *et al*., 2016; Rolando and Taylor, 2017). Whereas the requirements for Microprocessor binding and activity during microRNA biogenesis have been studied extensively and are known (Kim *et al*., 2017), Drosha-mediated cleavage of target mRNAs is an independent mechanism, and how sequence specificity is achieved to ensure physiological regulation of mRNAs is unclear (Kim *et al*., 2017; Rolando and Taylor, 2017). Furthermore, the binding of Drosha to an mRNA is not always conducive with cleavage, implying a multifaceted regulatory control of targeting and cleavage, and that these processes may not necessarily be linked (Rolando *et al*., 2016; Rolando and Taylor, 2017). Considering the ubiquitous expression of Drosha, we postulated that regulated target-specific activity of Drosha is controlled by the formation of specialized macromolecular complexes.

NFIB is a critical transcription factor in the regulation of glial cell differentiation. In the adult DG, NSCs retain tri-lineage potency but differentiation to oligodendrocytes is repressed by suppression of NFIB protein expression (Rolando *et al*., 2016). Drosha destabilizes NFIB mRNA in DG NSCs blocking gliogenesis and promoting neurogenesis. The NFIB mRNA contains two evolutionary conserved HPs located in the UTRs, and both are bound by Drosha, however, only the 3’ UTR HP is cleaved by Drosha (Rolando *et al*., 2016). We found that DGCR8 does not interact with NFIB UTR HPs, and its overexpression does not affect NFIB mRNA cleavage. Therefore, DGCR8 and the core Microprocessor are unlikely to contribute to Drosha-mediated regulation of NFIB in adult NSCs. This implies the requirement for other factors in cell-type specific regulation of NFIB mRNA stability.

Comparison of the MS^2^ data revealed major differences in the proteins associated with the 5’ UTR and 3’ UTR HPs suggesting different functions of these sequences in the control of NFIB expression. The NFIB 5’ UTR HP showed stronger association with ribosomal proteins including Rpl7, Rpl15, Rpl23a, Rpl27, Rpl31, Rps7, Rps15a and Rps25, than the NFIB 3’ UTR HP, suggesting a role in NFIB translation. This is supported by the GO analysis which revealed prominent roles of the 5’ UTR HP associated proteins in translational regulation. Whether Drosha plays a role in NFIB translational control through this 5’ UTR HP region remains unclear.

The protein with the highest fold enrichment on the NFIB 3’ UTR HP over the 5’ UTR HP was SAFB-like transcription modulator (Sltm), which together with Safb1 and Safb2 forms the Safb protein family. Sltm is expressed primarily in the nuclei and dendrites of cortical and hippocampal neurons where it may affect mRNA processing and/or transport (Norman et al., 2016). We also found that Stlm, Safb1 and Safb2 all bind to Drosha. As Safb proteins can form homo-and heterodimers, it is possible that they could form distinct multimeric complexes with Drosha that contribute to differences in function. Interestingly, although broadly expressed, Safb proteins are not functionally redundant as Safb1-deficiency results in a highly penetrant embryonic and perinatal lethality whereas Safb2-null mice are viable (Ivanova et al., 2005; Jiang et al., 2015). Unlike Safb1, Sltm and Safb2 do not bind selectively to the NFIB 3’ UTR HP. Therefore, it will be important to investigate the roles of Sltm and Safb2 in DG NSC fate regulation in order to assess the full regulatory repertoire of this family of RBPs. This is particularly interesting in the light of the finding that Safb2 also binds Drosha in human cells and assists in the processing of the non-optimal microRNA15a/16-1 pri-microRNA cluster (Hutter *et al*., 2020). Interestingly, Safb2 regulation of the Microprocessor on the microRNA15a/16-1 clustered does not require the Safb2 RNA-binding domain.

We identified Safb1 as a novel partner of Drosha in NSCs and a major regulator of Drosha-mediated NFIB mRNA cleavage. Safb1 binds AT-rich scaffold-matrix attachment regions (S/MARs) in DNA (Renz and Fackelmayer, 1996). However, Safb1 also binds RNA, recognizing GAA/AAG/AGA triplicates in a GAAGA consensus motif, and is involved in RNA-dependent chromatin organization and mRNA processing (Huo et al., 2020; Rivers *et al*., 2015; Stoilov et al., 2004; Van Nostrand *et al*., 2020). Due to the lethality of Safb1 knockout mice, the function of Safb1 in the brain was unclear (Ivanova *et al*., 2005). We show that Safb1 plays a surprising role in regulating cell fate in the adult hippocampus. Safb1 prevents oligodendrogenesis from tri-potent NSCs by activating Drosha-mediated cleavage of NFIB mRNA thereby preventing NFIB protein expression. This is in line with the Safb1 expression by adult DG NSCs, where oligodendrogenesis does not occur under physiological conditions, and lower expression by cells of the astrocytic and oligodendrocytic lineages and by oligodendrogenic NSCs of the postnatal V-SVZ (Rivers *et al*., 2015). In addition, reducing Safb1 levels curbed Drosha cleavage of NFIB mRNA in a dose-dependent fashion indicating that modulating Safb1 levels can regulate Drosha activity. In contrast to Safb2 regulation of the Microprocessor, Safb1 regulation of Drosha activity on the NFIB mRNA is independent of DGCR8, which does not bind the NFIB mRNA or affect NFIB mRNA stability (Rolando *et al*., 2016). Safb1 binds to the NFIB 3’ UTR HP but not to the NFIB 5’ UTR HP, indicating sequence specificity which this further supported by the mapping of Safb1 binding to NFIB mRNA in HepG cells by CLIP (Van Nostrand *et al*., 2020).

We show that Safb1 is necessary for Drosha cleavage of NFIB mRNA. Both the knockdown of Safb1 and expression of a mutant protein lacking the RE-domain with reduced Drosha binding ability, block Drosha cleavage of NFIB 3’ UTR HP. Although we did not formally show a requirement for Safb1-mediated RNA binding for NFIB mRNA processing, we identified multiple Safb1 binding motifs around the NFIB 3’ UTR HP that overlap with the Drosha cleavage sites and corelate with previous CLIP data showing Safb1 binding to the same region of NFIB mRNA (Rivers *et al*., 2015; Van Nostrand *et al*., 2020).

Although we focused on Safb1 here, we found many Drosha binding proteins that associated with the NFIB UTR HPs. Interestingly, Sam68, a member of the STAR family of RBPs, also affected NFIB HP processing. Sam68 has been shown to control RNA splicing including during neurogenesis (Chawla et al., 2009). Safb1 interacts with Sam68 and they co-distribute in the nucleus in response to stress (Sergeant et al., 2007). Sam68 has been shown to interact with the Microprocessor in different systems (Messina et al., 2012; Sellier et al., 2013). It is currently unclear whether Sam68 controls Drosha canonical or non-canonical functions. Qki is another member of the STAR protein family and isoform 5 of Qki is localized to the nucleus, the site of predominant Drosha expression. Of those RBPs we tested, Qki displayed the second strongest reduction in the expression of the NFIB reporter constructs. Qki is known to be involved in mRNA processing and splicing and in oligodendrocyte differentiation (Fagg et al., 2017; Larocque et al., 2005; Wu et al., 2002). Thus, Drosha may also contribute to the Qki regulation of oligodendrocyte formation and thus act at different levels in the lineage, however, this remains to be shown by future analyses. Although we focused here of RBPs that down regulated NFIB reporter expression, we found that Fus enhanced the levels of GFP from the NFIB 3’ UTR HP reporter. Interestingly, Fus also interacts with Safb1 (Yamaguchi and Takanashi, 2016). This raises the intriguing question of whether the collaboration between Fus and Safb1 extends beyond the DNA interactions previously reported and potentially to NFIB mRNA regulation.

Several mRNAs are now known to be targets of Drosha, and most of them contain evolutionarily conserved HP structures (Chong *et al*., 2010; Johanson et al., 2013; Knuckles *et al*., 2012; Rolando *et al*., 2016; Rolando and Taylor, 2017). More than 2000 human mRNAs are predicted to form secondary HPs that resemble pri-microRNAs, opening up a huge potential for non-canonical Drosha mRNA endonuclease function in different cells (Johanson *et al*., 2013; Pedersen et al., 2006).

Given the wide range of interacting proteins, it is conceivable that Drosha targets many more mRNAs than is currently appreciated and thus could control many biological processes. We propose that Drosha-mediated RNA cleavage requires interactions with specific RBPs that direct target specificity and activity. The differential targeting of Drosha to cell-type specific substrates has major implications in the dynamic control of Drosha activity and functions. In summary, our data uncovered novel Drosha interacting proteins and in the future it will be important to investigate these Drosha/RBP complexes and their functional involvement in NSC fate regulation as well as in other cellular systems and contexts.

## Materials and Methods

### Key resource table

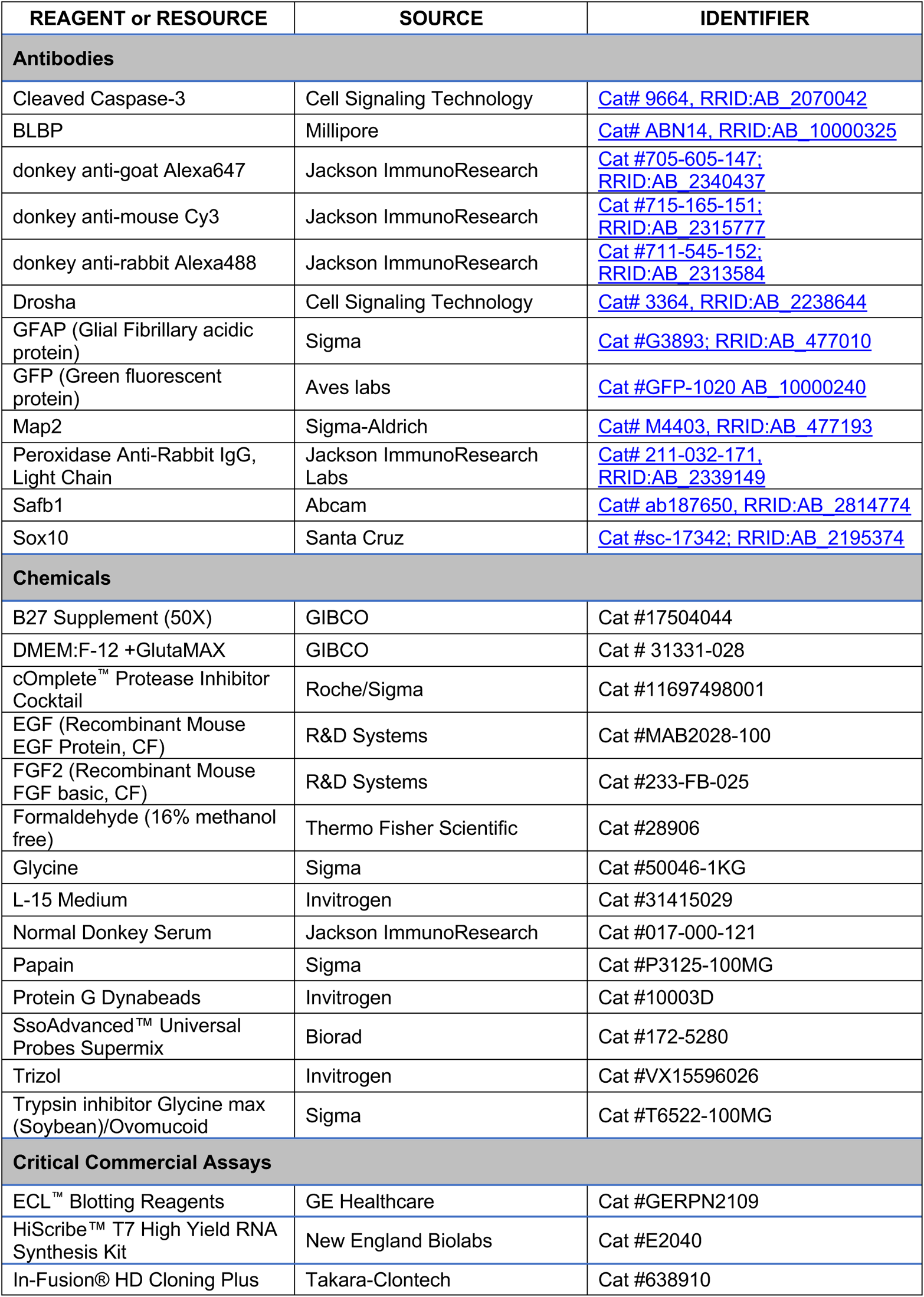
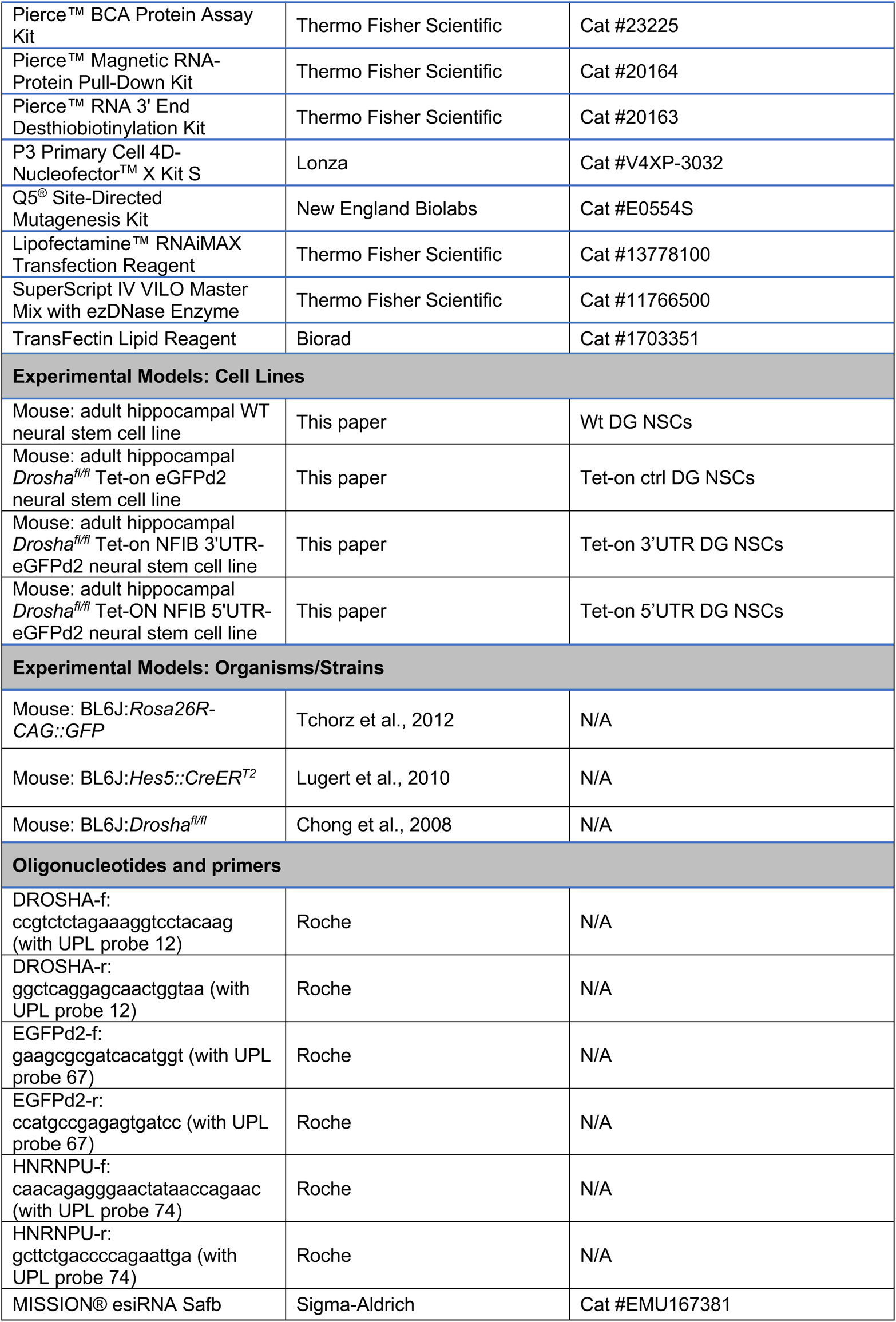
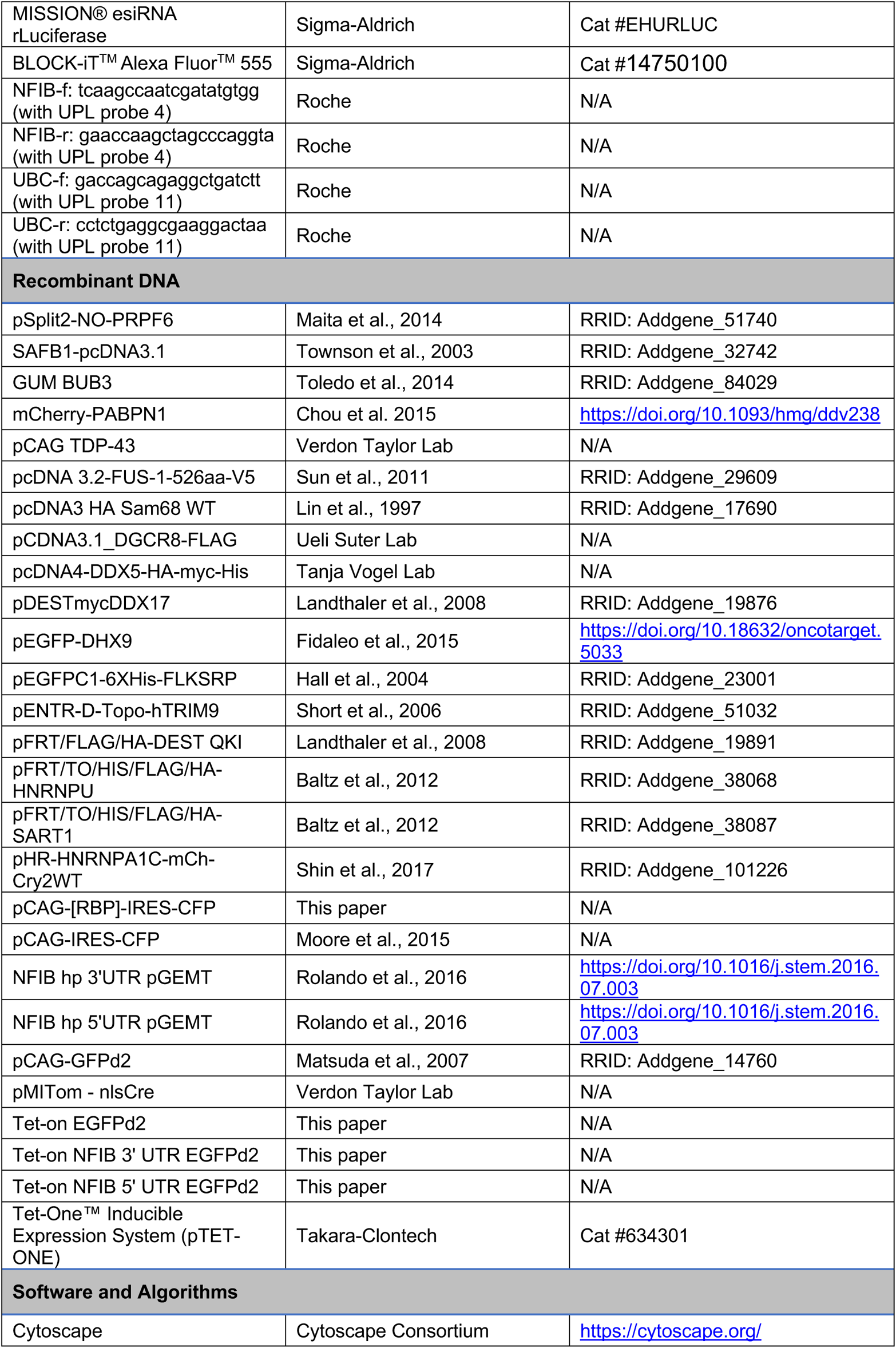
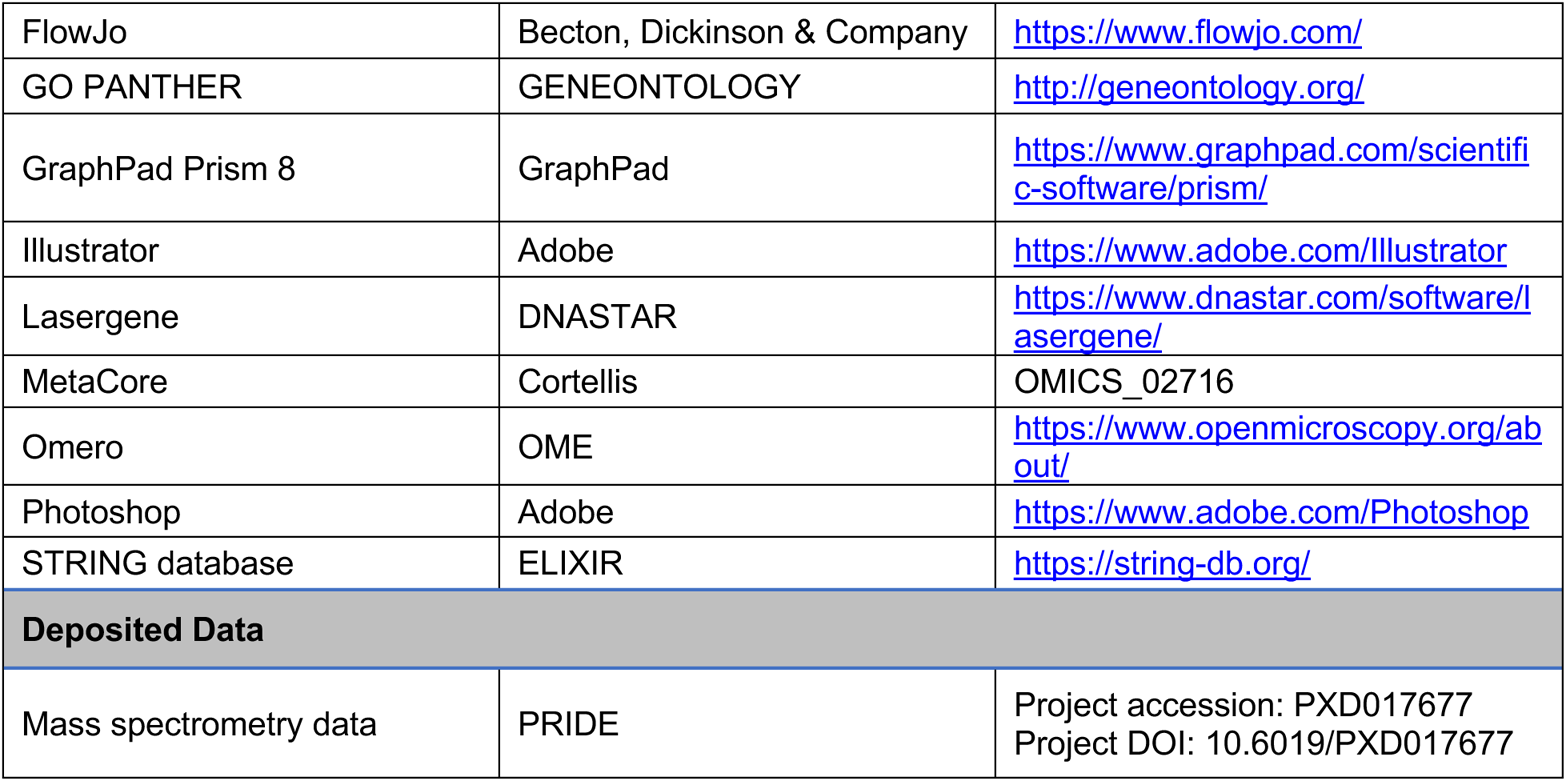

### Mouse strains and husbandry

The mouse models used in our experiments include: *Rosa26R-CAG::GFP*, *Hes5::CreER^T2^* (Lugert *et al*., 2010; Rolando *et al*., 2016) and *Drosha^fl/fl^* (Chong *et al*., 2010). No gender differences were observed. Mice were randomly selected for the experiments based on birth date and genotype. According to Swiss Federal and Swiss Veterinary office regulations, all mice were bred and kept in a specific pathogen-free animal facility with 12 hours day-night cycle and free access to clean food and water. All mice were healthy and immunocompetent. All procedures were approved by the Basel Cantonal Veterinary Office under license number 2537 (Ethics commission Basel-Stadt, Basel Switzerland).

### Hippocampal adult neural stem cell cultures

DG NSCs were isolated, cultured and differentiated, as described previously (Rolando *et al*., 2016; Zhang et al., 2019). 8-10 weeks-old mice were sacrificed in a CO_2_ chamber and decapitated. The brain was extracted, washed in ice cold sterile L15 medium (GIBCO) and live sectioned at 500 µm using a McIlwain’s tissue chopper. Brain slices were collected in cold HBSS, 10 mM HEPES and 100 I.U./mL penicillin and 100 μg/ml streptomycin in a 6 cm culture dish. After careful micro-dissection of the DG and removing the molecular layer and ventricular zone contaminants using a dissection binocular microscope, the dissected DGs were collected in cold HBSS, 10 mM HEPES and 100 I.U./mL penicillin and 100 μg/ml streptomycin in a 15 ml Falcon tube. After tissue sedimentation, the supernatant was removed and replaced by 100 µl pre-warmed Papain mix. The tissue pieces were incubated at 37°C in a water bath for 15 minutes with gentle agitation every 5 minutes, followed by addition of 50 µl of pre-warmed Trypsin inhibitor and incubation for 10 minutes at 37 °C. After adding 300 µl of DMEM/F12, the tissue was triturated with a 1 ml and 200 µl pipette tip. The sample was centrifuged at 80g to remove debris. The cell pellet was resuspended in DG NSC medium (DMEM:F12, Gibco, Invitrogen), 2% B27 (Gibco, Invitrogen), FGF2 20 ng/ml (R&D Systems), EGF 20 ng/ml (R&D Systems) and plated in a 48-well dish (Costar) coated with 100 μg/ml Poly-L-Lysine (Sigma) and 1 μg/ml Laminin (Sigma). Half of the cell medium was replaced every 2 days and cells were passaged every 6 days. Cells were passaged 6-20 times and characterized for marker expression and differentiation potential before use. Cell differentiation was induced by growth factor removal and continued culture for 6 days. Cell fixation for immunohistochemistry was performed for 10 minutes in 4% paraformaldehyde in 0.1 M phosphate buffer.

### Immunohistochemistry for brain tissues and NSC cultures

Mice were deeply anesthetized by injection of a ketamine/xylazine/acepromazine solution (150, 7.5 and 0.6 mg per kg body weight, respectively). Animals were perfused with ice-cold 0.9% saline followed by 4% paraformaldehyde in 0.1 M phosphate buffer. Brains were isolated and post-fixed overnight in 4% paraformaldehyde in 0.1 M phosphate buffer, and then cryoprotected with 30% sucrose in phosphate buffer at 4°C overnight. Brains were embedded and frozen in OCT (TissueTEK) and sectioned as 30 μm floating sections by cryostat (Leica). Free-floating coronal sections were stored at −20°C in antifreeze solution until use. DG NSCs fixation for immunohistochemistry was performed for 10 minutes in 4% paraformaldehyde in 0.1 M phosphate buffer.

Sections were incubated overnight at room temperature, with the primary antibody diluted in blocking solution of 1.5% normal donkey serum (Jackson ImmunoResearch), 0.5% Triton X-100 in phosphate-buffered saline. DG NSC cultures were incubate overnight at 4°C, with the primary antibody diluted in blocking solution of 1.5% normal donkey serum, 1% BSA, 0.2% Triton X-100 in phosphate-buffered saline.

Antibodies used: activated cleavedCASP3 (Cell Signaling, rabbit, 1:500), BLBP (Chemicon, rabbit, 1:400), GFP (AbD Serotec, sheep, 1:250; Invitrogen, rabbit, 1:700; AvesLabs, chicken, 1:500), Map2 (Sigma, mouse, 1:200), SAFB1 (Abcam, rabbit, 1:200), Sox10 (Santa Cruz, goat, 1:500) and GFAP (Sigma, mouse, 1:200). Sections were washed in phosphate-buffered saline and incubated at room temperature for 2 hours with the corresponding secondary antibodies in blocking solution. DG NSCs were washed in 1% BSA phosphate-buffered saline and incubated at room temperature for 35 minutes with the corresponding secondary antibodies in blocking solution.

Secondary antibodies and detection: Alexa488/Cy3/Alexa649 conjugated anti-chicken, mouse, goat, rabbit and sheep immunoglobulin (1:600, Jackson Immunoresearch). Sections were then washed and counter-stained with DAPI (1 μg/ml). Stained sections were mounted on Superfrost glass slides (Thermo Fisher Scientific), embedded in mounting medium containing diazabicyclo-octane (DABCO; Sigma) as an anti-fading agent. Brain sections and DG NSCs were visualized using a Zeiss LSM510 confocal microscope, Leica SP5 confocal microscope or Zeiss Apotome2 microscope. The antibody information is described in the KEY RESOURCES TABLE.

### Cell lysis for proteomics

DG NSCs were washed with PBS and incubated with lysis buffer (20 mM Hepes-KOH pH 7.9; 100 mM KCl; 0.2 mM EDTA; 0.2 mM PMSF, 1x complete proteinase inhibitor (Roche); 5% glycerol; 0.5% Triton) for 10 minutes on ice. Cells were scraped off with cell scraper and lysate was transferred in Eppendorf tube, followed by sonication for 5 cycles (Bioruptor, 30 seconds on/ 30 seconds off, 320W) at 4°C. Lysate was centrifuged at 13’000 x g for 10 minutes at 4°C. The supernatant was transferred in a new tube and BCA assay (Thermo Fischer Scientific, #23250) was performed according to the protocol of the manufacturer to determine protein concentration.

### Drosha co-immunoprecipitation assay

1 mg of total protein lysate was treated with or without RNAse, incubated with antibody (Drosha, Cell Signaling, 1:50 or Safb1, Abcam 1:50 resp.) on a rotating wheel overnight at 4°C (where 2.5% of each sample had been saved as input). Magnetic beads (Dynabeads, Thermo Fisher Scientific) were washed with 1 ml lysis buffer and DG cell lysate/Ab mix was added to the beads, followed by incubation for 4 hours at 4°C on a rotating wheel. Beads were washed 5 times with activated lysis buffer and resuspended in milli-Q water.

### Immunoblot analysis

5x Laemmli Buffer was added to the samples to reach a final volume of 1x and heated 5 minutes at 95 °C at 700 rpm. Protein samples were separated on 10% SDS-polyacrylamide gels and transferred to PVDF membrane (Immobilon). The membrane was blocked for 1h with 5% BSA in TBS-T followed by overnight incubation at 4°C with antibody (Drosha 1:1000; D28B1, Cell Signaling, in 5 %BSA; Safb1, 1:1000 ab187650, 2.5% BSA; PKC-alpha [H-7] (SC-8393) 1:500 1% BSA). Membrane was washed 3 times for 5 minutes with TBS-T followed by secondary antibody incubation (rabbit anti light chain HRP 211-032-171, 1:5000) for 1h at room temperature. Membrane was washed 3 times with TBS-T and once with TBS. Bands were detected by chemiluminescence (ECL, GE Healthcare).

### Affinity purification and sample preparation for MS-based proteome analysis

IP and pull-down probes of DG NSCs were subjected to on-bead digestion (Hubner et al., 2010) by trypsin (5 µg/ml, Promega) in 1.6 M Urea / 0.1 M Ammonium bicarbonate buffer at 27 C for 30 minutes. Supernatant eluates containing active trypsin were further incubated with 1 mM TCEP at room temperature overnight. Carbamidomethylation of cysteine residues was performed next using 5 mM Chloroacetamide in the dark for 30 minutes. The tryptic digest was acidified (pH<3) using TFA and desalted using C18 reversed phase spin columns (Harvard Apparatus) according to the protocol of the manufacturer. Dried peptides were dissolved in 0.1% aqueous formic acid solution at a concentration of 0.2 mg/ml prior to injection into the mass spectrometer.

### Mass spectrometry analysis

For each sample, aliquots of 0.4 μg of total peptides were subjected to LC-MS analysis using a dual pressure LTQ-Orbitrap Elite mass spectrometer connected to an electrospray ion source (both Thermo Fisher Scientific) and a custom-made column heater set to 60°C. Peptide separation was carried out using an EASY nLC-1000 system (Thermo Fisher Scientific) equipped with a RP-HPLC column (75μm × 30cm) packed in-house with C18 resin (ReproSil-Pur C18–AQ, 1.9 μm resin; Dr. Maisch GmbH, Germany) using a linear gradient from 95% solvent A (0.1% formic acid in water) and 5% solvent B (80% acetonitrile, 0.1% formic acid, in water) to 35% solvent B over 50 minutes to 50% solvent B over 10 minutes to 95% solvent B over 2 minutes and 95% solvent B over 18 minutes at a flow rate of 0.2 μl/min. The data acquisition mode was set to obtain one high resolution MS scan in the FT part of the mass spectrometer at a resolution of 240,000 full width at half maximum (at 400 m/z, MS1) followed by MS/MS (MS^2^) scans in the linear ion trap of the 20 most intense MS signals. The charged state screening modus was enabled to exclude unassigned and singly charged ions and the dynamic exclusion duration was set to 30 seconds. The collision energy was set to 35%, and one microscan was acquired for each spectrum. The mass spectrometry proteomics data in this manuscript have been deposited to the ProteomeXchange Consortium via the PRIDE (Perez-Riverol et al., 2019) partner repository with the accession number: PXD017677 and project DOI: 10.6019/PXD017677.

### Protein identification and label-free quantification

The acquired raw-files were imported into the Progenesis QI software (v2.0, Nonlinear Dynamics Limited), which was used to extract peptide precursor ion intensities across all samples applying the default parameters. The generated mgf-files were searched using MASCOT against a decoy database containing normal and reverse sequences of the *Mus musculus* proteome (UniProt, April 2017) and commonly observed contaminants (in total 34490 sequences) generated using the SequenceReverser tool from the MaxQuant software (Version 1.0.13.13). The following search criteria were used: full tryptic specificity was required (cleavage after lysine or arginine residues, unless followed by proline); 3 missed cleavages were allowed; carbamidomethylation (C) was set as fixed modification; oxidation (M) and protein N-terminal acetylation were applied as variable modifications; mass tolerance of 10 ppm (precursor) and 0.6 Da (fragments). The database search results were filtered using the ion score to set the false discovery rate (FDR) to 1% on the peptide and protein level, respectively, based on the number of reverse protein sequence hits in the datasets.

Quantitative analysis results from label-free quantification were normalized and statically analyzed using the SafeQuant R package v.2.3.4 (https://github.com/eahrne/SafeQuant/) (Ahrne et al., 2016) to obtain protein relative abundances. This analysis included global data normalization by equalizing the total peak/reporter areas across all LC-MS runs, summation of peak areas per protein and LC-MS/MS run, followed by calculation of protein abundance ratios. Only isoform specific peptide ion signals were considered for quantification. The summarized protein expression values were used for statistical testing of between condition differentially abundant proteins. Here, empirical Bayes moderated t-Tests were applied, as implemented in the R/Bioconductor limma package (http://bioconductor.org/packages/release/bioc/html/limma.html). The resulting per protein and condition comparison P-values were adjusted for multiple testing using the Benjamini-Hochberg method. All LC-MS analysis runs are acquired from independent biological samples. To meet additional assumptions (normality and homoscedasticity) underlying the use of linear regression models and Student t-Test MS-intensity signals are transformed from the linear to the log-scale. Unless stated otherwise linear regression was performed using the ordinary least square (OLS) method as implemented in *base* package of R v.3.1.2 (http://www.R-project.org/).

The sample size of three biological replicates was chosen assuming a within-group MS-signal Coefficient of Variation of 10%. When applying a two-sample, two-sided Student t-test this gives adequate power (80%) to detect protein abundance fold changes higher than 1.65, per statistical test. Note that the statistical package used to assess protein abundance changes, SafeQuant, employs a moderated t-Test, which has been shown to provide higher power than the Student’s t-test. We did not do any simulations to assess power, upon correction for multiple testing (Benjamini-Hochberg correction), as a function of different effect sizes and assumed proportions of differentially abundant proteins.

### RNA biotinylation

NFIB 3’ UTR and 5’ UTR HP forming regions were *in vitro* transcribed using T7 transcriptase (NEB, E2040) and purified with Trizol extraction (described in RT-qPCR paragraph). RNA was biotinylated using the Pierce^TM^ RNA 3’ End Desthiobiotinylation Kit (Thermo Fisher Scientific). 50 pmol of RNA was incubated in T4 RNA ligase buffer at 16 °C overnight. RNA ligase was extracted with 100 µl chloroform:isoamyl alcohol (Sigma, C0549) and centrifuged 2 minutes at maximum speed. Aqueous phase was precipitated overnight in 5M NaCl. Sample was centrifuged at 13’000 x g for 20 minutes at 4 °C. Pellet was washed with 70% ethanol and resuspended in dH_2_O. Labeling efficiency was determined by dot blotting using the chemiluminescent nucleic acid detection module kit (89880, Thermo Fisher scientific).

### RNA pull-down of NSC proteins

Experiment was performed as described in the manufacturers protocol (Pierce^TM^ Magnetic RNA-Protein Pull-Down Kit, Thermo Fisher Scientific, 20164). 50 µl of streptavidin magnetic beads were washed 3 times with 20 mM Tris, followed by 1x RNA capture buffer. 50 pmol of previously desthiobiotinylated RNA was added to the beads and incubated for 30 minutes on a heated shaker plate at 24 °C and 750 rpm.

Beads were washed 3 times with 20 mM Tris and 100 µl 1x Protein-RNA Binding Buffer, followed by 100 µl of RNA Protein binding buffer including 60 µl of Protein lysate (2mg/ml) (Cell lysis described in corresponding paragraph) for 60 minutes 4 °C with agitation. The beads were washed 3 times with 100 µl of the kit 1x wash buffer and then further processed for WB or MS^2^.

### Enrichment analysis and candidate selection

Datasets of significantly enriched proteins were analyzed for process networks by MetaCore (Cortellis) and molecular processes by GO terms (PANTHER, GENEONTOLOGY). An interaction network was drawn using STRING database (ELIXIR). Percental enrichment in MetaCore bar plot was calculated as listed in corresponding category out of total dataset. Significance is determined by P-value. For STRING network analysis, only experimentally determined data and curated databases were considered. Nodes with ≥ 1 Edge are displayed. Network was visualized and analyzed with Cytoscape software (Shannon et al., 2003). Prime candidates for functional analysis in reporter assay were selected using the following criteria: (i) enrichment in MS datasets; (ii) relevance in MetaCore, GO and STRING analysis; (iii) Drosha interactions reported in the literature

### Stable NSC line generation

NFIB 3’ UTR, 5’ UTR and EGFPd2 sequences were cloned in the multiple cloning site of pTet-One vector (Takara-Clontech, 634301). DG NSC *Drosha^fl/fl^* cells were brought in suspension by incubating with 0.25% trypsin (Gibco #15090) in Versene (Gibco #15040) for 5 minutes at 37°C followed by centrifugation at 80g for 5 minutes at RT. Cells were electroporated with the corresponding DNA vector using a 4D-Nucleofector (Lonza, program DS-112) according to protocol of manufacturer and re-plated in plastic dish (Costar) coated with 100 μg/ml Poly-L-Lysine (Sigma) and 1 μg/ml Laminin (Sigma). Cells were induced with 1 μg/ml doxycycline in DG NSC medium. 48 hours post induction, cells were sorted for GFP^+^ by flow cytometry (FACSaria III, BD Biosciences). All GFP^+^ cells were collected, centrifuged at 80g for 5 minutes and resuspended in DG NSC medium. Cells were re-plated, and passaged 3 times. Thereafter, cells were re-induced with 1 μg/ml doxycycline and re-sorted two more times. Correct genotype of cell line was confirmed by genotyping using NFIB UTR and EGFPd2 specific primers.

### RBP overexpression vector generation

Coding sequences of selected prime candidates were cloned in a *CAG-IRES-CFP* expression vector, using the In-Fusion cloning kit (Takara) following manufactures protocols. Protein cDNAs from source vector were amplified by PCR and cloned upstream of IRES-CFP element of target vector. Sequences of all resulting vectors were verified by Sanger-Sequencing (Eurofins).

### Tet-on DG NSC nucleofection and FACS and analysis

Tet-on ctrl., Tet-on 3’ UTR HP and 5’ UTR HP DG NSCs were brought in suspension by incubating with 0.25% trypsin (Gibco #15090) in Versene (Gibco #15040) for 4 minutes at 37°C followed by centrifugation at 80g for 5 minutes at RT. Cells were electroporated with either CFP-IRES or RBP-IRES-CFP vector (5 µg) using a 4D-Nucleofector (Lonza, program DS-112) in 16-well stripes (500,000 cells/well) and re-plated in plastic dish (Costar) coated with 100 μg/ml Poly-L-Lysine (Sigma) and 1 μg/ml Laminin (Sigma). For rescue experiments, Tet-on ctrl., Tet-on 3’ UTR HP and 5’ UTR HP *Drosha^fl/fl^* DG NSCs were electroporated with 1.6 µg of *Cre-IRES-Tomato* and 3.4 µg of *CAG::IRES-CFP* or *CAG::Safb-IRES-CFP* with 4D-Nucleofector (program DS-112).

After 24 hours or 48 hours cells were induced with 1 μg/ml doxycycline in DG NSC medium. 48 hours post-induction, cells were brought in suspension with 0.25% trypsin in Versene collected in DMEM/F12 w/o red phenol, filtered through a 40 μm cell sieve (Miltenyi Biotec) and analyzed by flow cytometry (FACSaria III, BD Biosciences). GFP and CFP double negative, GFP-single positive and CFP-single positive DG NSCs were used to create the compensation matrix and set the sorting gates. At least 100,000 events were recorded for each sample and MFI (median fluorescence intensity) of the population of interest was subsequently analyzed with FlowJO (Becton Dickinson). GFP^+^, CFP^+^GFP^+^, Tom^+^ cells were sorted, centrifuged at 80g for 5 minutes and used for RNA isolation and gene expression analysis (see below).

### Safb1-ΔRE-HA mutant generation and overexpression

The Safb1-ΔRE-HA mutant was generated using the Q5 Site-Directed Mutagenesis Kit (NEB). 5’-overlapping primers were designed and the Safb1-HA vector was amplified excluding the RE region (amino acids 552-708). The vector was re-ligated, transformed and endotoxin free maxiprep (Qiagen) DNA sequenced. The Safb1-ΔRE mutant variant was cloned into the same CAG-IRES-CFP expression used to express the RBPs in the RBP overexpression analysis (see RBP overexpression vector generation). 2 million N2A cells were transfected using TransFectin (Biorad) with either CFP-ctrl., Safb1-HA-IRES-CFP or Safb1-ΔRE-HA-IRES-CFP / CFP-ctrl (1:5) (10ug total in all samples). 48 hours after transfection the cells were prepared for co-IP experiments (see Cell Lysis and Drosha co-Immunoprecipitation assay). Tet-on ctrl. and Tet-on 3’ UTR HP DG NSCs were electroporated with either CFP-ctrl., Safb1-HA-IRES-CFP or Safb1-ΔRE-HA-IRES-CFP and analysed by FACS as described above (see Tet-on DG NSC nucleofection and FACS and analysis).

### Cross-linking immunoprecipitation (CLIP)

WT mouse DG NSCs in culture were washed with cold PBS and crosslinked at 254 nm, 300 mJ/cm^2^ in a BioLink UV-Crosslinker. Cells were lysed with RIPA buffer (0.1 M sodium phosphate pH 7.2, 150 mM sodium chloride, 0.1% SDS, 1% sodium deoxycholate, 1% NP-40, 1 mM activated Na_3_VO_4_, 1 mM NaF, 1x complete protease inhibitor (Roche)). Lysate was collected and centrifuged for 5 minutes at 13’000 RPM at 4°C. Sample was treated with 10 µl DNase (Roche) and incubated for 5 minutes in a heated shaker at 37°C and 1000 RPM, followed by centrifugation at full speed at 4°C and Input probe separation. 50 µl magnetic beads (Dynabeads, Invitrogen) were washed with lysis buffer and incubated with target specific antibody (anti-Safb1, Abcam) for 1 hour at RT on a rotating wheel. Beads were washed twice with lysis buffer and incubated with cross-linked NSC lysate for 2 hours at 4°C on a rotating wheel. Beads were washed 5 times with lysis buffer, incubated with 4 mg/ml Proteinase K (Roche) in PK buffer (100 mM Tris HCL pH7.5, 50 mM NaCl, 10 mM EDTA) for 20 minutes at 37°C at 1000 RPM, followed by downstream RNA analysis. PCR primers for the genes of interest were designed by the Universal Probe Library Assay Design Center (Roche) (primer sequences listed in key resource table).

### RNA isolation and RT-qPCR

Total RNA was isolated following the standard phenol-chloroform protocol from the manufacturer (Trizol, Life Technologies). 100% Trizol was added to the sample, followed by 20% of total volume chloroform. Sample was centrifuged at 12’000 x g for 15 minutes at 4°C. Aqueous phase was extracted and RNA was precipitated with isopropanol and washed with ethanol. RNA pellet was resuspended in RNase-free Milli-Q water. Reverse transcription was performed using Superscript IV Vilo following manufacturers protocol (Thermo Fischer Scientific, 11766050). RNA was incubated for 5 minutes with ezDNAse enzyme at 37°C, followed by incubation with SuperScript™ IV VILO™ Master Mix at 25°C for 10 minutes, 50°C for 10 minutes and 85°C for 5 minutes. For expression analysis of genes of interest, we used the comparative Ct method using Rpl13a and Actin as normalizing genes. Experiments were performed using a qTOWER^3^ real-time PCR machine (Analytik Jena). Three biological replicates for each genotype and three technical replicates for each gene were analyzed.

### esiRNA-mediated Safb1 RNA knock-down

WT DG NSCs were brought in suspension with 0.25% trypsin in Versene followed by centrifugation at 80g for 5 minutes at RT. Cells were electroporated with either 70 pmol esiRNA Luciferase or esiRNA Safb1 (Sigma) in combination with 30 pmol RNA probe Alexa555 (Sigma) using a 4D-Nucleofector (Lonza, program DS-112) in 16-well stripes (500,000 cells/well) and re-plated on glass coverslips coated with 100 μg/ml Poly-L-Lysine (Sigma) and 1 μg/ml Laminin (Sigma). Cells were fixed 48, 72 and 96 hours after nucleofection in 4% paraformaldehyde in 0.1 M phosphate buffer for 10 minutes. N2A cells were transfected with Lipofectamine RNAiMAX (Thermo Fisher Scientific) according to manufacturer’s instructions. Cells were transfected with 90 pmol of esiRNA Luciferase or esiRNA Safb1 (Sigma) in a 6-well plate, seeded one day before transfection (250’000 cells/well in DMEM 4.5g/l, 10%FBS). 48 hours after transfection, cells were lysed RIPA lysis buffer for downstream analysis, followed by sonication for 5 cycles (Bioruptor, 30 seconds on/ 30 seconds off, 320W) at 4°C. Lysates were centrifuged at 13’000 x g for 10 minutes at 4°C. The supernatants were transferred to a new tube and protein quantified by BCA assay (Thermo Fischer Scientific, #23250) according to the manufacturer instructions.

### Analysis of Safb1 on Drosha activity by *in vitro* processing assay

N2A cells were transfected with esiRNA Safb1 as described above (esiRNA-mediated Safb1 RNA knock-down). After 48h, samples were lysed and the full lysate was treated with RNAse A and DNAse I and immunoprecipitated for Drosha as described above (see Cell Lysis and Drosha co-Immunoprecipitation assay). For the *in vitro* processing assay (Rolando *et al*., 2016), samples were mixed in the following ratios: Sample 1 (15 ul WT), Sample 2 (10 ul WT, 5 ul KD), Sample 3 (5 ul WT, 10 ul KD), Sample 4 (15 ul KD). A total of 30 μl of the processing reaction were prepared and contained: 15 μl of beads from Drosha immunoprecipitated beads, mixed or control fraction, 6.4 mM MgCl2, 0.75 μl RNase Inhibitor (Invitrogen) and 0.5 μg RNA probe containing T7 RNA polymerase (NEB) transcribed NFIB 3’ UTR HP RNA or AR RNA / NFIB 3’ UTR RNA hybrid RNA as control. The reaction was carried out at 25°C for 30 minutes. RNA was extracted using Trizol and subsequently analyzed by RT qPCR.

### GFP reporter analysis after glial differentiation

Tet-on ctrl and Tet-on 3’ UTR HP DG NSCs were cultured under expansion conditions until 70-80% confluent and then astrocytic differentiation induced by removal of growth factors EGF and FGF and treatment with 10% fetal bovine serum (FBS). After 5 days, the cells were induced with 1 μg/ml doxycycline. 48 hours post-doxycycline induction, the cells were trypsinized with 0.25% trypsin in Versene, collected in DMEM/F12 w/o red phenol, stained with BV421 labelled anti-CD44 or BV421 labelled rat IgG2b Isotype control in 0.05% BSA for 20 min on ice protected from light, filtered through a 40 μm cell sieve (Miltenyi Biotec) and analyzed by flow cytometry (FACS Aria III, BD Biosciences). GFP and CD44 double-negative, GFP-single positive and CD44-single positive NSCs were used to create the compensation matrix and set the sorting gates. At least 100,000 events were recorded for each sample and MFI (median fluorescence intensity) of the population of interest was subsequently analyzed with FlowJO (Becton Dickinson).

### Safb1 overexpression in postnatal neural stem cell neurosphere cultures culture

Postnatal C57BL/6 neurospheres were prepared as described previously (Giachino et al., 2009). Postnatal mice were sacrificed, decapitated and their brains collected in 6 cm dishes containing sterile cold HBSS. The meninges were carefully removed and the striatum isolated by dissection. The lateral walls of the stiatum were dissected and the tissue was dissociated in 500 µl pre-warmed Papain solution at 37°C for 30 minutes followed by addition of 500 µl of pre-warmed Trypsin inhibitor and incubation for a further 5 minutes at 37 °C. The tissue was mechanically dissociated with 1 ml and 200 µl pipette tips. After addition of 9 ml of DMEM/F12, the samples were centrifuged at 80g for 5 minutes to remove debris. The cell pellet from each animal was resuspended in neurosphere medium DMEM:F12 (Gibco, Invitrogen), 2% B27 (Gibco, Invitrogen), EGF 10 ng/ml (R&D Systems) and plated in T25 flasks. The cells were fed every 2 days with fresh neurosphere medium. The cells were passaged every 4 days.

Dissociated neurosphere cells were electroporated with either CFP-IRES or Safb-IRES-CFP vector (4µg) using a 4D-Nucleofector (Lonza, program DS-112) in 16-well stripes (500,000 cells/well) and re-plated onto 13 mm glass cover slips in 24 well plates coated with 100 μg/ml Poly-L-Lysine (Sigma) and 1 μg/ml Laminin (Sigma) in neurosphere medium. Two days later, differentiation was induced by replacing the medium with neurosphere medium without EGF. The cells were differentiated for 2 days and then fixed for immunocytochemical analysis by 10 minutes incubation in 4% paraformaldehyde in 0.1 M phosphate buffer. The cells were then stained using antibodies against anti-GFP (Chicken, Aves labs, 1:500), anti-Safb1 (Rabbit, Abcam, 1:300) and anti-Sox10 (Goat, Santa Cruz, 1:200) antibodies (see Key Resources Table). The experiment was repeated three times with 3 biological replicates each.

### Quantification and statistical analysis

Images of immunostainings were captured and processed on a Confocal Leica SP5 and Apotome2 (Zeiss). According to the Swiss governmental guidelines and requirements, the principles of the 3Rs for animal research were taking into consideration to reduce the number of mice used in the experiment. Data are presented as averages of indicated number of samples. Data representation and statistical analysis were performed using GraphPad Prism software. Statistical comparisons were conducted by two-tailed unpaired Student’s t test, Mann-Whitney test or one-way ANOVA test, as indicated. The size of samples (n) is described in the figure legends. Statistical significance is determined by p values (*p <0.05, **p <0.01, ***p < 0.001) and error bars are presented as SEM.

For Safb1 motif analysis, the top 20 of pentamers (ranked for the highest Z-scores) identified in previous Safb1 iCLIP experiments (Rivers *et al*., 2015) were mapped on the NFIB 3’UTR sequence. These most enriched pentamers were also used to create the consensus binding motif with the web-based WebLogo software (http://weblogo.berkeley.edu/).

## Author Contributions

N.I., C.R., E.-A.B., P.F., and T.M. designed and performed experiments and evaluated and interpreted the data. T.B. performed the MS^2^ analysis and interpreted the data. V.T. conceived and designed the project and evaluated and interpreted the data. N.I., C.R., and V.T. wrote the paper and prepared the figures. V.T. acquired the financing for the project. All authors edited the manuscript.

## Acknowledgments

We thank the members of the V.T. laboratory for helpful discussions and Frank Sager for excellent technical assistance. We thank the Proteomics Facility of the Biozentrum Basel and the Animal Core Facility of the University of Basel. This work was supported by the Swiss National Science Foundation (31003A_162609 and 31003A_182388).

## References

1. Ahrne, E., Glatter, T., Vigano, C., Schubert, C., Nigg, E.A., and Schmidt, A. (2016). Evaluation and Improvement of Quantification Accuracy in Isobaric Mass Tag-Based Protein Quantification Experiments. J Proteome Res 15, 2537–2547. 10.1021/acs.jproteome.6b00066.

2. Altmeyer, M., Toledo, L., Gudjonsson, T., Grofte, M., Rask, M.B., Lukas, C., Akimov, V., Blagoev, B., Bartek, J., and Lukas, J. (2013). The chromatin scaffold protein SAFB1 renders chromatin permissive for DNA damage signaling. Mol Cell 52, 206–220. 10.1016/j.molcel.2013.08.025.

3. Baser, A., Skabkin, M., Kleber, S., Dang, Y., Gulculer Balta, G.S., Kalamakis, G., Gopferich, M., Ibanez, D.C., Schefzik, R., Lopez, A.S., et al. (2019). Onset of differentiation is post-transcriptionally controlled in adult neural stem cells. Nature 566, 100–104. 10.1038/s41586-019-0888-x.

4. Beckervordersandforth, R., and Rolando, C. (2019). Untangling human neurogenesis to understand and counteract brain disorders. Curr Opin Pharmacol 50, 67–73. 10.1016/j.coph.2019.12.002.

5. Berg, D.A., Su, Y., Jimenez-Cyrus, D., Patel, A., Huang, N., Morizet, D., Lee, S., Shah, R., Ringeling, F.R., Jain, R., et al. (2019). A Common Embryonic Origin of Stem Cells Drives Developmental and Adult Neurogenesis. Cell 177, 654–668 e615. 10.1016/j.cell.2019.02.010.

6. Boldrini, M., Fulmore, C.A., Tartt, A.N., Simeon, L.R., Pavlova, I., Poposka, V., Rosoklija, G.B., Stankov, A., Arango, V., Dwork, A.J., et al. (2018). Human Hippocampal Neurogenesis Persists throughout Aging. Cell Stem Cell 22, 589–599 e585. 10.1016/j.stem.2018.03.015.

7. Bonaguidi, M.A., Wheeler, M.A., Shapiro, J.S., Stadel, R.P., Sun, G.J., Ming, G.L., and Song, H. (2011). In vivo clonal analysis reveals self-renewing and multipotent adult neural stem cell characteristics. Cell 145, 1142–1155. 10.1016/j.cell.2011.05.024.

8. Bonzano, S., Crisci, I., Podlesny-Drabiniok, A., Rolando, C., Krezel, W., Studer, M., and De Marchis, S. (2018). Neuron-Astroglia Cell Fate Decision in the Adult Mouse Hippocampal Neurogenic Niche Is Cell-Intrinsically Controlled by COUP-TFI In Vivo. Cell Rep 24, 329–341. 10.1016/j.celrep.2018.06.044.

9. Chawla, G., Lin, C.H., Han, A., Shiue, L., Ares, M., Jr., and Black, D.L. (2009). Sam68 regulates a set of alternatively spliced exons during neurogenesis. Mol Cell Biol 29, 201–213. 10.1128/MCB.01349-08.

10. Chong, M.M., Zhang, G., Cheloufi, S., Neubert, T.A., Hannon, G.J., and Littman, D.R. (2010). Canonical and alternate functions of the microRNA biogenesis machinery. Genes Dev 24, 1951–1960. 10.1101/gad.1953310.

11. Deneen, B., Ho, R., Lukaszewicz, A., Hochstim, C.J., Gronostajski, R.M., and Anderson, D.J. (2006). The transcription factor NFIA controls the onset of gliogenesis in the developing spinal cord. Neuron 52, 953–968. 10.1016/j.neuron.2006.11.019.

12. Eriksson, P.S., Perfilieva, E., Bjork-Eriksson, T., Alborn, A.M., Nordborg, C., Peterson, D.A., and Gage, F.H. (1998). Neurogenesis in the adult human hippocampus. Nat Med 4, 1313–1317. 10.1038/3305.

13. Fagg, W.S., Liu, N., Fair, J.H., Shiue, L., Katzman, S., Donohue, J.P., and Ares, M., Jr. (2017). Autogenous cross-regulation of Quaking mRNA processing and translation balances Quaking functions in splicing and translation. Genes Dev 31, 1894–1909. 10.1101/gad.302059.117.

14. Gage, F.H. (2019). Adult neurogenesis in mammals. Science 364, 827–828. 10.1126/science.aav6885.

15. Gerstberger, S., Hafner, M., and Tuschl, T. (2014). A census of human RNA-binding proteins. Nat Rev Genet 15, 829–845. 10.1038/nrg3813.

16. Giachino, C., Basak, O., and Taylor, V. (2009). Isolation and manipulation of mammalian neural stem cells in vitro. Methods Mol Biol 482, 143–158. 10.1007/978-1-59745-060-7_9.

17. Goncalves, J.T., Schafer, S.T., and Gage, F.H. (2016). Adult Neurogenesis in the Hippocampus: From Stem Cells to Behavior. Cell 167, 897–914. 10.1016/j.cell.2016.10.021.

18. Han, J., Lee, Y., Yeom, K.H., Kim, Y.K., Jin, H., and Kim, V.N. (2004). The Drosha-DGCR8 complex in primary microRNA processing. Genes Dev 18, 3016–3027. 10.1101/gad.1262504.

19. Han, J., Pedersen, J.S., Kwon, S.C., Belair, C.D., Kim, Y.K., Yeom, K.H., Yang, W.Y., Haussler, D., Blelloch, R., and Kim, V.N. (2009). Posttranscriptional crossregulation between Drosha and DGCR8. Cell 136, 75–84. 10.1016/j.cell.2008.10.053.

20. Huang, R., Han, M., Meng, L., and Chen, X. (2018). Transcriptome-wide discovery of coding and noncoding RNA-binding proteins. Proc Natl Acad Sci U S A 115, E3879–E3887. 10.1073/pnas.1718406115.

21. Hubner, N.C., Bird, A.W., Cox, J., Splettstoesser, B., Bandilla, P., Poser, I., Hyman, A., and Mann, M. (2010). Quantitative proteomics combined with BAC TransgeneOmics reveals in vivo protein interactions. J Cell Biol 189, 739–754. 10.1083/jcb.200911091.

22. Huo, X., Ji, L., Zhang, Y., Lv, P., Cao, X., Wang, Q., Yan, Z., Dong, S., Du, D., Zhang, F., et al. (2020). The Nuclear Matrix Protein SAFB Cooperates with Major Satellite RNAs to Stabilize Heterochromatin Architecture Partially through Phase Separation. Mol Cell 77, 368–383 e367. 10.1016/j.molcel.2019.10.001.

23. Hutter, K., Lohmuller, M., Jukic, A., Eichin, F., Avci, S., Labi, V., Szabo, T.G., Hoser, S.M., Huttenhofer, A., Villunger, A., and Herzog, S. (2020). SAFB2 Enables the Processing of Suboptimal Stem-Loop Structures in Clustered Primary miRNA Transcripts. Mol Cell 78, 876–889 e876. 10.1016/j.molcel.2020.05.011.

24. Ivanova, M., Dobrzycka, K.M., Jiang, S., Michaelis, K., Meyer, R., Kang, K., Adkins, B., Barski, O.A., Zubairy, S., Divisova, J., et al. (2005). Scaffold attachment factor B1 functions in development, growth, and reproduction. Mol Cell Biol 25, 2995–3006. 10.1128/MCB.25.8.2995-3006.2005.

25. Jiang, S., Katz, T.A., Garee, J.P., DeMayo, F.J., Lee, A.V., and Oesterreich, S. (2015). Scaffold attachment factor B2 (SAFB2)-null mice reveal non-redundant functions of SAFB2 compared with its paralog, SAFB1. Dis Model Mech 8, 1121–1127. 10.1242/dmm.019885.

26. Johanson, T.M., Lew, A.M., and Chong, M.M. (2013). MicroRNA-independent roles of the RNase III enzymes Drosha and Dicer. Open Biol 3, 130144. 10.1098/rsob.130144.

27. Kang, W., Nguyen, K.C.Q., and Hebert, J.M. (2019). Transient Redirection of SVZ Stem Cells to Oligodendrogenesis by FGFR3 Activation Promotes Remyelination. Stem Cell Reports 12, 1223–1231. 10.1016/j.stemcr.2019.05.006.

28. Kempermann, G., Gage, F.H., Aigner, L., Song, H., Curtis, M.A., Thuret, S., Kuhn, H.G., Jessberger, S., Frankland, P.W., Cameron, H.A., et al. (2018). Human Adult Neurogenesis: Evidence and Remaining Questions. Cell Stem Cell 23, 25–30. 10.1016/j.stem.2018.04.004.

29. Kim, B., Jeong, K., and Kim, V.N. (2017). Genome-wide Mapping of DROSHA Cleavage Sites on Primary MicroRNAs and Noncanonical Substrates. Mol Cell 66, 258–269 e255. 10.1016/j.molcel.2017.03.013.

30. Knuckles, P., Vogt, M.A., Lugert, S., Milo, M., Chong, M.M., Hautbergue, G.M., Wilson, S.A., Littman, D.R., and Taylor, V. (2012). Drosha regulates neurogenesis by controlling neurogenin 2 expression independent of microRNAs. Nat Neurosci 15, 962–969. 10.1038/nn.3139.

31. Lachapelle, F., Avellana-Adalid, V., Nait-Oumesmar, B., and Baron-Van Evercooren, A. (2002). Fibroblast growth factor-2 (FGF-2) and platelet-derived growth factor AB (PDGF AB) promote adult SVZ-derived oligodendrogenesis in vivo. Mol Cell Neurosci 20, 390–403. 10.1006/mcne.2002.1124.

32. Larocque, D., Galarneau, A., Liu, H.N., Scott, M., Almazan, G., and Richard, S. (2005). Protection of p27(Kip1) mRNA by quaking RNA binding proteins promotes oligodendrocyte differentiation. Nat Neurosci 8, 27–33. 10.1038/nn1359.

33. Lee, D., and Shin, C. (2018). Emerging roles of DROSHA beyond primary microRNA processing. RNA Biol 15, 186–193. 10.1080/15476286.2017.1405210.

34. Li, X., Zhao, X., Fang, Y., Jiang, X., Duong, T., Fan, C., Huang, C.C., and Kain, S.R. (1998). Generation of destabilized green fluorescent protein as a transcription reporter. J Biol Chem 273, 34970–34975. 10.1074/jbc.273.52.34970.

35. Lugert, S., Basak, O., Knuckles, P., Haussler, U., Fabel, K., Gotz, M., Haas, C.A., Kempermann, G., Taylor, V., and Giachino, C. (2010). Quiescent and active hippocampal neural stem cells with distinct morphologies respond selectively to physiological and pathological stimuli and aging. Cell Stem Cell 6, 445–456. 10.1016/j.stem.2010.03.017.

36. Lugert, S., Vogt, M., Tchorz, J.S., Muller, M., Giachino, C., and Taylor, V. (2012). Homeostatic neurogenesis in the adult hippocampus does not involve amplification of Ascl1(high) intermediate progenitors. Nat Commun 3, 670. 10.1038/ncomms1670.

37. Macias, S., Cordiner, R.A., Gautier, P., Plass, M., and Caceres, J.F. (2015). DGCR8 Acts as an Adaptor for the Exosome Complex to Degrade Double-Stranded Structured RNAs. Mol Cell 60, 873–885. 10.1016/j.molcel.2015.11.011.

38. Messina, V., Meikar, O., Paronetto, M.P., Calabretta, S., Geremia, R., Kotaja, N., and Sette, C. (2012). The RNA binding protein SAM68 transiently localizes in the chromatoid body of male germ cells and influences expression of select microRNAs. PLoS One 7, e39729. 10.1371/journal.pone.0039729.

39. Moreno-Jimenez, E.P., Flor-Garcia, M., Terreros-Roncal, J., Rabano, A., Cafini, F., Pallas-Bazarra, N., Avila, J., and Llorens-Martin, M. (2019). Adult hippocampal neurogenesis is abundant in neurologically healthy subjects and drops sharply in patients with Alzheimer’s disease. Nat Med 25, 554–560. 10.1038/s41591-019-0375-9.

40. Nguyen, T.A., Jo, M.H., Choi, Y.G., Park, J., Kwon, S.C., Hohng, S., Kim, V.N., and Woo, J.S. (2015). Functional Anatomy of the Human Microprocessor. Cell 161, 1374–1387. 10.1016/j.cell.2015.05.010.

41. Norman, M., Rivers, C., Lee, Y.B., Idris, J., and Uney, J. (2016). The increasing diversity of functions attributed to the SAFB family of RNA-/DNA-binding proteins. Biochem J 473, 4271–4288. 10.1042/BCJ20160649.

42. Obernier, K., and Alvarez-Buylla, A. (2019). Neural stem cells: origin, heterogeneity and regulation in the adult mammalian brain. Development 146, dev156059. 10.1242/dev.156059.

43. Pedersen, J.S., Bejerano, G., Siepel, A., Rosenbloom, K., Lindblad-Toh, K., Lander, E.S., Kent, J., Miller, W., and Haussler, D. (2006). Identification and classification of conserved RNA secondary structures in the human genome. PLoS Comput Biol 2, e33. 10.1371/journal.pcbi.0020033.

44. Perez-Riverol, Y., Csordas, A., Bai, J., Bernal-Llinares, M., Hewapathirana, S., Kundu, D.J., Inuganti, A., Griss, J., Mayer, G., Eisenacher, M., et al. (2019). The PRIDE database and related tools and resources in 2019: improving support for quantification data. Nucleic Acids Res 47, D442–D450. 10.1093/nar/gky1106.

45. Pilaz, L.J., and Silver, D.L. (2015). Post-transcriptional regulation in corticogenesis: how RNA-binding proteins help build the brain. Wiley Interdiscip Rev RNA 6, 501–515. 10.1002/wrna.1289.

46. Pilz, G.A., Bottes, S., Betizeau, M., Jorg, D.J., Carta, S., Simons, B.D., Helmchen, F., and Jessberger, S. (2018). Live imaging of neurogenesis in the adult mouse hippocampus. Science 359, 658–662. 10.1126/science.aao5056.

47. Ratti, A., Fallini, C., Cova, L., Fantozzi, R., Calzarossa, C., Zennaro, E., Pascale, A., Quattrone, A., and Silani, V. (2006). A role for the ELAV RNA-binding proteins in neural stem cells: stabilization of Msi1 mRNA. J Cell Sci 119, 1442–1452. 10.1242/jcs.02852.

48. Renz, A., and Fackelmayer, F.O. (1996). Purification and molecular cloning of the scaffold attachment factor B (SAF-B), a novel human nuclear protein that specifically binds to S/MAR-DNA. Nucleic Acids Res 24, 843–849. 10.1093/nar/24.5.843.

49. Rivers, C., Idris, J., Scott, H., Rogers, M., Lee, Y.B., Gaunt, J., Phylactou, L., Curk, T., Campbell, C., Ule, J., et al. (2015). iCLIP identifies novel roles for SAFB1 in regulating RNA processing and neuronal function. BMC Biol 13, 111. 10.1186/s12915-015-0220-7.

50. Rolando, C., Erni, A., Grison, A., Beattie, R., Engler, A., Gokhale, P.J., Milo, M., Wegleiter, T., Jessberger, S., and Taylor, V. (2016). Multipotency of Adult Hippocampal NSCs In Vivo Is Restricted by Drosha/NFIB. Cell Stem Cell 19, 653–662. 10.1016/j.stem.2016.07.003.

51. Rolando, C., and Taylor, V. (2017). Non-canonical post-transcriptional RNA regulation of neural stem cell potential. Brain Plast 3, 111–116. 10.3233/BPL-170046.

52. Rouillard, A.D., Gundersen, G.W., Fernandez, N.F., Wang, Z., Monteiro, C.D., McDermott, M.G., and Ma’ayan, A. (2016). The harmonizome: a collection of processed datasets gathered to serve and mine knowledge about genes and proteins. Database (Oxford) 2016. 10.1093/database/baw100.

53. Sellier, C., Freyermuth, F., Tabet, R., Tran, T., He, F., Ruffenach, F., Alunni, V., Moine, H., Thibault, C., Page, A., et al. (2013). Sequestration of DROSHA and DGCR8 by expanded CGG RNA repeats alters microRNA processing in fragile X-associated tremor/ataxia syndrome. Cell Rep 3, 869–880. 10.1016/j.celrep.2013.02.004.

54. Sergeant, K.A., Bourgeois, C.F., Dalgliesh, C., Venables, J.P., Stevenin, J., and Elliott, D.J. (2007). Alternative RNA splicing complexes containing the scaffold attachment factor SAFB2. J Cell Sci 120, 309–319. 10.1242/jcs.03344.

55. Seri, B., Garcia-Verdugo, J.M., Collado-Morente, L., McEwen, B.S., and Alvarez-Buylla, A. (2004). Cell types, lineage, and architecture of the germinal zone in the adult dentate gyrus. J Comp Neurol 478, 359–378. 10.1002/cne.20288.

56. Shannon, P., Markiel, A., Ozier, O., Baliga, N.S., Wang, J.T., Ramage, D., Amin, N., Schwikowski, B., and Ideker, T. (2003). Cytoscape: a software environment for integrated models of biomolecular interaction networks. Genome Res 13, 2498–2504. 10.1101/gr.1239303.

57. Sohn, J., Selvaraj, V., Wakayama, K., Orosco, L., Lee, E., Crawford, S.E., Guo, F., Lang, J., Horiuchi, M., Zarbalis, K., et al. (2012). PEDF is a novel oligodendrogenic morphogen acting on the adult SVZ and corpus callosum. J Neurosci 32, 12152–12164. 10.1523/JNEUROSCI.0628-12.2012.

58. Sorrells, S.F., Paredes, M.F., Cebrian-Silla, A., Sandoval, K., Qi, D., Kelley, K.W., James, D., Mayer, S., Chang, J., Auguste, K.I., et al. (2018). Human hippocampal neurogenesis drops sharply in children to undetectable levels in adults. Nature 555, 377–381. 10.1038/nature25975.

59. Spadotto, V., Giambruno, R., Massignani, E., Mihailovich, M., Maniaci, M., Patuzzo, F., Ghini, F., Nicassio, F., and Bonaldi, T. (2020). PRMT1-mediated methylation of the microprocessor-associated proteins regulates microRNA biogenesis. Nucleic Acids Res 48, 96–115. 10.1093/nar/gkz1051.

60. Spalding, K.L., Bergmann, O., Alkass, K., Bernard, S., Salehpour, M., Huttner, H.B., Bostrom, E., Westerlund, I., Vial, C., Buchholz, B.A., et al. (2013). Dynamics of hippocampal neurogenesis in adult humans. Cell 153, 1219–1227. 10.1016/j.cell.2013.05.002.

61. Stoilov, P., Daoud, R., Nayler, O., and Stamm, S. (2004). Human tra2-beta1 autoregulates its protein concentration by influencing alternative splicing of its pre-mRNA. Hum Mol Genet 13, 509–524. 10.1093/hmg/ddh051.

62. Szklarczyk, D., Gable, A.L., Lyon, D., Junge, A., Wyder, S., Huerta-Cepas, J., Simonovic, M., Doncheva, N.T., Morris, J.H., Bork, P., et al. (2019). STRING v11: protein-protein association networks with increased coverage, supporting functional discovery in genome-wide experimental datasets. Nucleic Acids Res 47, D607–D613. 10.1093/nar/gky1131.

63. Tobin, M.K., Musaraca, K., Disouky, A., Shetti, A., Bheri, A., Honer, W.G., Kim, N., Dawe, R.J., Bennett, D.A., Arfanakis, K., and Lazarov, O. (2019). Human Hippocampal Neurogenesis Persists in Aged Adults and Alzheimer’s Disease Patients. Cell Stem Cell 24, 974–982 e973. 10.1016/j.stem.2019.05.003.

64. Townson, S.M., Dobrzycka, K.M., Lee, A.V., Air, M., Deng, W., Kang, K., Jiang, S., Kioka, N., Michaelis, K., and Oesterreich, S. (2003). SAFB2, a new scaffold attachment factor homolog and estrogen receptor corepressor. J Biol Chem 278, 20059–20068. 10.1074/jbc.M212988200.

65. Townson, S.M., Sullivan, T., Zhang, Q.P., Clark, G.M., Osborne, C.K., Lee, A.V., and Oesterreich, S. (2000). HET/SAF-B overexpression causes growth arrest and multinuclearity and is associated with aneuploidy in human breast cancer. Clinical Cancer Research 6, 3788–3796.

66. Treiber, T., Treiber, N., Plessmann, U., Harlander, S., Daiss, J.L., Eichner, N., Lehmann, G., Schall, K., Urlaub, H., and Meister, G. (2017). A Compendium of RNA-Binding Proteins that Regulate MicroRNA Biogenesis. Mol Cell 66, 270–284 e213. 10.1016/j.molcel.2017.03.014.

67. Van Nostrand, E.L., Pratt, G.A., Yee, B.A., Wheeler, E.C., Blue, S.M., Mueller, J., Park, S.S., Garcia, K.E., Gelboin-Burkhart, C., Nguyen, T.B., et al. (2020). Principles of RNA processing from analysis of enhanced CLIP maps for 150 RNA binding proteins. Genome Biol 21, 90. 10.1186/s13059-020-01982-9.

68. Wu, J.I., Reed, R.B., Grabowski, P.J., and Artzt, K. (2002). Function of quaking in myelination: regulation of alternative splicing. Proc Natl Acad Sci U S A 99, 4233–4238. 10.1073/pnas.072090399.

69. Yamaguchi, A., and Takanashi, K. (2016). FUS interacts with nuclear matrix-associated protein SAFB1 as well as Matrin3 to regulate splicing and ligand-mediated transcription. Sci Rep 6, 35195. 10.1038/srep35195.

70. Yoo, A.S., Sun, A.X., Li, L., Shcheglovitov, A., Portmann, T., Li, Y., Lee-Messer, C., Dolmetsch, R.E., Tsien, R.W., and Crabtree, G.R. (2011). MicroRNA-mediated conversion of human fibroblasts to neurons. Nature 476, 228–231. 10.1038/nature10323.

71. Zhang, R., Boareto, M., Engler, A., Louvi, A., Giachino, C., Iber, D., and Taylor, V. (2019). Id4 Downstream of Notch2 Maintains Neural Stem Cell Quiescence in the Adult Hippocampus. Cell Rep 28, 1485–1498 e1486. 10.1016/j.celrep.2019.07.014.

